# New insights into the role of *Cutibacterium acnes*-derived extracellular vesicles in inflammatory skin disorders

**DOI:** 10.1101/2022.12.15.520547

**Authors:** Maria Pol Cros, Júlia Mir-Pedrol, Lorena Toloza, Nastassia Knödlseder, Marc Güell, Julien Maruotti, Christos C. Zouboulis, Maria-José Fábrega Fernández

## Abstract

**Background:** *Cutibacterium acnes* (*C. acnes*) is one of the most prevalent bacteria that form the human skin microbiota and, depending on multifactorial conditions it can help to maintain the skin homeostasis. Actually, different phylotypes of *C. acnes* have been associated with different degrees of acne vulgaris development, while others, such as the H1 subtype, have been detected in patients with non-acneic skin. However, due to the physiology of the skin, the skin microbiota neither has direct access to the skin’s sebaceous glands nor to the main immune cells, as they are protected by a sebum layer. Therefore, the inter-kingdom communication relies on secreted factors and bacterial extracellular vesicles (EVs). In this context, the purpose of this project was to study the role of EVs secreted by three different phylotypes of *C. acnes* (A1 as pathogenic, H1 as beneficial and H2 as commensal).

**Results:** Main findings showed that the proteomic profile of the cargo embodied in the EVs reflects unique characteristics of the different *C. acnes* phylotypes in terms of lifestyle, survival and virulence. Moreover, *in vitro* skin models showed an extended pro-inflammatory modulation of A1 EVs, while H1 EVs displayed a high sebum-reducing potential.

**Conclusions:** This study has highlighted the role of *C. acnes* EVs as key modulators during skin alterations, specially H1 EVs as an alternative based-natural treatment to fight acne vulgaris symptomatology.

## 1. INTRODUCTION

The human body harbours a flourishing diversity of microbes that form a dynamic and functional system that develops in synergy with the physiological development of its host (Rackaityte & Lynch, 2020). The human microbiome resides primarily in the skin, the oral mucosa and the gastrointestinal tracts (Paetzold et al., 2019). Recent data demonstrates that the specific composition of the microbial community is associated with health and disease, which suggests that a detailed characterization of the microbial community would reveal important commensal host-microbiome as well as microbe-microbe interactions with therapeutic implications (Blum, 2017). Over the past decade, researchers have discovered evidence of extensive communication between bacteria, skin cells and immune cells. These interactions are aimed at strengthening and repairing the barrier formed by the skin, reinforcing the body’s defences against infection and reducing excess inflammation (Eisenstein, 2020). Nonetheless, failure of this communication can disrupt the protective barrier of the skin and lead to dysbiosis between commensals and pathogens.

One of the most common bacteria present in the normal human skin microbiota is the Gram-positive anaerobic bacterium, *Cutibacterium acnes* (*C. acnes*) which dominates the upper 1.9mm of terminal of pilosebaceous units (Jahns & Alexeyev, 2014; McLaughlin et al., 2019). *C. acnes* helps to preserve and support the natural microbial balance of the skin but under certain conditions, it can also substantially alter its local environment and cause disease (Christensen et al., 2016; O’Neill & Gallo, 2018; Shu et al., 2013). In fact, this bacterium is linked to a wide range of skin diseases, including acne vulgaris. Although the aetiology and pathogenesis of acne are still uncertain, microbiome alterations are thought to be one of the principal mechanisms behind the development of acne (Paetzold et al., 2019). Recently, some authors have described that different phylotypes of *C. acnes* have different degrees of association with acne vulgaris (Fitz-Gibbon et al., 2013; Lomholt & Kilian, 2010; McDowell et al., 2013; McLaughlin et al., 2019). This brings into question how bacteria communicate with host cells to maintain homeostasis of the skin microbiota if sebaceous glands and immune cells are protected by a layer of sebum from external environment and skin microorganisms. This communication is known to be mediated by bacterial extracellular vesicles (EVs), which are nano-sized lipid bilayer vesicles approximately 20-500 nm in diameter that can pass through the lipid barrier and interact with host cells (Bajic et al., 2020; Brown et al., 2015).

Since EVs play a key role in interkingdom communication and immune modulation, in the present study, different *in vitro* models mimicking human skin have been established to assay the potential of different *C. acnes* EVs. In this context and based on previous publications (Choi et al., 2018a; Chudzik et al., 2022; Paetzold et al., 2019), we decided to study the role that EVs from three different phylotypes of *C. acnes* play in the onset of acne, considering *C. acnes* A1 phylotype (clade IA1) as a pathogenic phylotype, *C. acnes* H1 phylotype (clade IB) as a probiotic phylotype, and *C. acnes* H2 phylotype (clade IB) as a commensal one. Moreover, proteomic characterization of *C. acnes* A1, H1 and H2 EVs has been carried out to determine the relationship of their protein cargo with the skin microbiota. Finally, a novel model mimicking acne-prone skin has been set up in which the sebum-reducing potential of EVs from different *C. acnes* phylotypes has also been tested.

## 2. MATERIALS AND METHODS

### 2.1 Bacterial phylotypes and culture conditions

*C. acnes* phylotypes used in this study were extracted from healthy human skin (A1 and H1) or commercially acquired (H2, KPA17202 from DSMZ), typed by SLST typing and we previously sequenced. Sequencing data can be found in the European nucleotide Archive under project number PRJEB42527.

Initial cultures were always started from glycerol stocks (stored at −80°C) and seeded on Brucella agar plates which were incubated for 4-5 days at 37°C under anaerobic conditions using the BD GasPak EZ anaerobic bag system.

### 2.2 Isolation of EVs

The process followed for the isolation of EVs consisted of several steps: cultivation, centrifugation, concentration or ultrafiltration and ultracentrifugation (Fig. S1).

In the cultivation step, *C. acnes* phylotypes A1, H1 and H2 were collected from a Brucella agar plate using sterile swabs (Deltalab) and were transferred into 1 L of Brucella liquid medium. After 7 to 10 days at 37°C with 300 rpm of stirring and maintaining anaerobic conditions, the culture was centrifuged in 250 mL sterile bottles at 10,000 x g for 30 min and at 4°C. For this step, a Beckman Coulter Avanti JXN-26 high-velocity centrifuge with the J-LITE JLA-16.250 Fixed-Angle Aluminium Rotor were used. Then, the pellet was discarded, and the supernatant (SN) was filtered using the Thermo Scientific Nalgene Rapid-Flow 75 mm Filter Unit (500 mL) to avoid bacterial contamination. Afterwards, the SN was concentrated by a Centricon Plus-70 10 KDa system using the JS-5.3 Swinging-Bucket Rotor. To ensure free contamination during the process, an extra step of 0.22 μm filtration was added to the protocol. The filtered and concentrated SN was then ultra-centrifuged in a Beckman Coulter optima L-100 XP ultracentrifuge using the SW41 Ti Swinging-Bucket Rotor at 154,300 x g for 2h and at 4°C to precipitate the EVs. The pellet containing the isolated *C. acnes* EVs was washed with 1X phosphate buffered-saline (PBS) buffer and ultra-centrifuged again for 1h under the same conditions as in the previous step.

Finally, the SN was discarded, and the pellet was resuspended with 200 μL of PBS buffer and kept in an Eppendorf at −20°C. To ensure non-bacterial presence in EVs samples, 10 μL of each batch were seeded in a Brucella agar plate and incubated for 4-5 days at 37°C under anaerobic conditions.

### 2.3 EVs labelling protocol

EVs labelling was performed as previously described by (Bajic et al., 2020; Cañas et al., 2016; Fábrega et al., 2017). Briefly, isolated A1 EVs were resuspended in PBS buffer, centrifuged at 154,300 x g for 2h at 4°C, and resuspended in labelling buffer (50 mM Na2CO3, 100 mM NaCl, pH 9.2) in the presence of 1 mg/mL of octadecyl rhodamine B-R18 (Invitrogen) and incubated for 1 h at 25°C. Labelled A1 EVs were pelleted by ultracentrifugation at 154,300 x g for 1h at 4°C, resuspended in PBS buffer (0.2 M NaCl) and washed three times to fully remove the unbound dye. After the final ultracentrifugation step, B-R18 labelled A1 EVs were resuspended in PBS buffer (0.2 M NaCl) containing a protease inhibitory cocktail (Complete Protease Inhibitory Tablets, Roche) and were stored at −20°C.

### 2.4 Negative staining and Transmission Electron Microscopy (TEM)

Isolated *C. acnes* H1 EVs were examined by TEM after negative staining, as described in previous publications (Aguilera et al., 2014). In this particular case, EVs were resuspended in TRIS buffer as PBS buffer is not compatible with uranyl acetate. A drop of EVs suspension was absorbed for 2 min on Formvar/carbon-coated grids that were previously activated with UV light. The grids were washed with distilled water, stained with 2% uranyl acetate for 1 min, air dried and examined by TEM (*Parc Científic de Barcelona*).

### 2.5 Nanoparticle Tracking Analysis (NTA) of EVs

EVs resuspended in 200 μL of PBS buffer were sent to *Institut de Ciència de materials de Barcelona* to be examined with the Malvern Nanosight NS300 instrument. This system uses NTA technology characterize nanoparticles in suspension in the size range of 10-2000 nm. A video is taken and the NTA software tracks the Brownian motion of individual EVs and calculates their size and total concentration.

### 2.6 Determination of protein concentration and SDS-PAGE

Protein concentrations in EVs samples were determined using the Qubit Protein Assay Kit (Thermo Fisher Scientific). Measuring was done using the Qubit Fluorometer (Thermo Fisher Scientific). This assay was performed at room temperature using specific Qubit assay tubes (Cat. no. Q32856).

For protein separation, the samples were subjected to sodium dodecyl sulfate-polyacrylamide gel electrophoresis (SDS-PAGE) using the Mini Gel Tank electrophoresis system (Invitrogen). Proteins were denatured in a loading buffer (4X LDS sample buffer +β-mercaptoethanol) at 98°C for 5 min. Then, 10 μL of PageRuler Protein Ladder and 7 μg of each sample of proteins mixed with 5 μL of loading buffer were loaded and separated on a NuPAGE 4 to 12% Bis-Tris Gel (Invitrogen) using MOPS SDS Running Buffer 20X (Life technologies) at a constant voltage of 100 V and for 40 min. After the run was completed, several bands were identified. Gel images were visualized in the ChemiDoc MP Imaging System (Bio-Rad).

### 2.7 Proteomic analysis: EVs preparation

EVs were lysed to get a total representation of their protein content. To do that, liquid nitrogen (N2) was used to obtain beads of the EVs solutions of each *C. acnes* phylotype. Then, 0.5 g of beads were disrupted using the Freezer Mill. Conditions used for the lysis were: precooling for 5 min, running step for 2 min, 2 cycles of interval and at a rate of 10 cycles per second (CPS). The powder obtained was resuspended in RIPA buffer supplemented with complete protease inhibitory tablets (Roche). Afterwards, total protein was precipitated with acetone and the final precipitate was resuspended with a quantity ranging from 50 μL to 250 μL of 6 M urea to have a minimum of 8 μg of protein. Qubit protein assay was performed to determine the estimated quantity of protein in each EV sample.

### 2.8 Identification of EVs proteins by LC-MS/MS analysis

For the identification of the EVs proteins, mass spectrometry was performed by the UPF/CRG Proteomics Unit.

Samples (8 μg) were reduced with dithiothreitol (24 nmol, 37°C, 60 min) and alkylated in the dark with iodoacetamide (48 nmol, 25°C, 30 min). The resulting protein extract was first diluted to 2M urea with 200 mM ammonium bicarbonate for digestion with endoproteinase Lys-C (1:10 w:w, 37°C, o/n, Wako, cat # 129-02541), and then diluted 2-fold with 200 mM ammonium bicarbonate for trypsin digestion (1:10 w:w, 37°C, 8h, Promega cat # V5113). After digestion, the peptide mix was acidified with formic acid and desalted with a MicroSpin C18 column (The Nest Group, Inc) prior to LC-MS/MS analysis.

Afterwards, samples were analysed using the LTQ - Orbitrap Fusion Lumos mass spectrometer (Thermo Fisher Scientific, San Jose, CA, USA) coupled with an EASY-nLC 1200 (Thermo Fisher Scientific (Proxeon), Odense, Denmark). Peptides were loaded directly onto the analytical column and were separated by reversed-phase chromatography using a 50-cm column with an inner diameter of 75 μm, packed with a 2 μm C18 particles spectrometer (Thermo Scientific).

Chromatographic gradients started at 95% buffer A and 5% buffer B with a flow rate of 300 nL/min for 5 min and gradually increased to 25% buffer B and 75% A in 79 min and then to 40% buffer B and 60% A in 11 min. After each analysis, the column was washed for 10 min with 10% buffer A and 90% buffer B. Buffer A: 0.1% formic acid in water. Buffer B: 0.1% formic acid in 80% acetonitrile.

The mass spectrometer was operated in positive ionization mode with nanospray voltage set at 2.4 kV and source temperature at 305°C. The acquisition was performed in data-dependent acquisition (DDA) mode and full MS scans with 1 micro scan at a resolution of 120,000 were used over a mass range of m/z 350-1400 with detection in the Orbitrap mass analyser, auto gain control (AGC) was set to “Standard” and maximum injection time to “Auto”. In each cycle of data-dependent acquisition analysis, following each survey scan, the most intense ions above a threshold ion count of 10000 were selected for fragmentation. The number of selected precursor ions for fragmentation was determined by the “Top Speed” acquisition algorithm and a dynamic exclusion of 60 seconds. Fragment ion spectra were produced via high-energy collision dissociation (HCD) at a normalized collision energy of 28% and they were acquired in the ion trap mass analyser. AGC was set to 2E4, and an isolation window of 0.7 m/z and a maximum injection time of 12 ms were used. Digested bovine serum albumin (New England Biolabs cat # P8108S) was analysed between each sample to avoid sample carryover and to assure stability of the instrument and QCloud (Chiva et al., 2018) has been used to control instrument longitudinal performance during the project.

Acquired spectra were analysed using the Proteome Discoverer software suite (v2.0, Thermo Fisher Scientific) and the Mascot search engine (v2.6, Matrix Science (Perkins et al., n.d.)). The data were searched against NCBI *acnes* phylotype H2171202 database (as in August 2021, 2476 entries, https://www.ncbi.nlm.nih.gov/nuccore/NC_006085.1). For peptide identification a precursor ion mass tolerance of 7 ppm was used for MS1 level, trypsin was chosen as the enzyme, and up to three missed cleavages were allowed. The fragment ion mass tolerance was set to 0.5 Da for MS2 spectra. Oxidation of methionine and N-terminal protein acetylation were used as variable modifications whereas carbamidomethylation on cysteines was set as a fixed modification. The false discovery rate (FDR) in peptide identification was set to a maximum of 5%. Peptide quantification data were retrieved from the “Precursor ion area detector” node from Proteome Discoverer (v2.0) using 2 ppm mass tolerance for the peptide extracted ion current (XIC). The obtained values were used to calculate protein fold changes and their corresponding adjusted p values.

### 2.9 Bioinformatic analysis

Proteins found by mass spectrometry were annotated from the UniProt database (UniProt Consortium, 2021). Pseudo proteins or proteins lacking information in UniProt were removed from the analysis. Only proteins with an area found in two or three of the replicas were considered. The mean of the areas of each protein was calculated.

Proteins present only in one of the three phylotype EVs were selected for further analysis. The information on biological processes, cellular components and molecular functions was obtained and manually curated to classify each protein into a category.

### 2.10 *In vitro* human cell lines maintenance

The immortalized keratinocytes cell line (HaCaT), bought in the American Type Culture Collection (ATCC), were cultured in T-75 culture flasks (Thermo Scientific) with Dulbecco’s Modified Eagle Medium (DMEM, Gibco) supplemented with 10% of Fetal Bovine Serum (FBS) and 0.1% of penicillin and streptomycin. They were incubated at 37°C in 5% of CO2, and 95% of humidity until 90-100% of confluency.

Immortalized human SZ95 sebocytes (Zouboulis et al., 1999) were cultured in T-75 culture flasks (Thermo Scientific) with Sebomed basal medium (DMEM/F12, Gibco) supplemented with 10% of FBS, 0.1% of penicillin and streptomycin, 5 ng/mL of recombinant human epidermal growth factor (EGFr) and 500 μL of filtrated calcium chloride (CaCl2) 1M. They were incubated at 37°C in 5% of CO2, and 95% of humidity until 90-100% of confluency.

The immortalized T lymphocytes cell line (Jurkat, Clone E6-1), bought in the ATCC, were cultured in T-75 culture flasks (Thermo Scientific) with Roswell Park Memorial Institute medium (RPMI, Gibco) supplemented with 10% of FBS and 0.1 % of penicillin and streptomycin. They were incubated at 37°C in 5% of CO2, and 95% of humidity until 90-100% of confluency.

PCi-SEB, derived from human iPSC with a Caucasian phototype (Phenocell), were seeded at a density of 25.000 viable cells/cm^2^ on fibronectin-coated tissue culture plate (Falcon) in PhenoCULT-SEB basal medium supplemented with 0.1% of supplement A. Then, the media was changed and supplemented with 0.1% of supplement M and incubated for 2 days more until cells reached 90% of confluence and they were differentiated into mature sebocytes. Cells were incubated at 37°C in 5% CO2 and 95% of humidity during the whole differentiation process.

### 2.11 Internalization of labelled-EVs

To monitor EVs uptake by both HaCaT and SZ95 sebocytes, a total of 1·10^5^ cells were seeded in an 8-well chamber slider (Ibidi) until approximately 80% of confluence. Before the assay, the medium was aspirated and replaced with rhodamine B-R18-labelled A1 EVs (2 μg/mL) suspended in DMEM and Sebomed both without fetal bovine serum and red phenol. The cells with the labelled A1 EVs were incubated at 37°C in 5% of CO2, and 95% of humidity for 24h.

### 2.12 Confocal fluorescence microscopy

Cells were fixed for 30 min with 3% of paraformaldehyde at room temperature (RT) and washed with PBS buffer to eliminate possible EVs residues. Plasma membranes were labelled with fluorescent Alexa-488 wheat germ agglutinin (WGA) (Invitrogen) and nuclei with Hoechst (Invitrogen). For this, cells were incubated for 25 min with Alexa-488 WGA (1 μg/mL) and during 10-15 min with Hoechst (3 drops/mL). After PBS buffer washing, Ibidi mounting medium was added (3 drops/well) to prepare the cells for microscopic visualisation.

Confocal microscopy was carried out using a ZEISS LSM 980 with Airyscan 2 confocal microscope, using the 63x oil immersion objective lens. Fluorescence was recorded at 405 nm (blue; Hoechst), 488 nm (green; WGA) and 546 nm (red; rhodamine B-R18). Z-stack images were taken at 1.0-μm. Images were analysed using the Fiji image processing package.

### 2.13 Cytotoxicity assay

To evaluate cytotoxicity, an XTT Cell Viability Assay (Invitrogen) was performed following the manufacturer’s protocol. The assay kit includes the XTT reagent and an Electron Coupling Reagent. In this case, the XTT reagent, is sensitive to cellular redox potential and in the presence of live cells converts from a water-soluble compound to an orange-coloured formazan product. 2· 10^5^ cells/mL of HaCaT and SZ95 sebocytes were seeded in two different 96-well plates. After 24h, cells were incubated with 1.56 μg/mL, 3.125 μg/mL, 6.25 μg/mL, 12.5 μg/mL, 25 μg/mL, 50 μg/mL, 100 μg/mL and 200 μg/mL of *C. acnes* A1, H1 and H2 EVs for 24h at 37°C in 5% of CO2 and 95% of humidity. Then, 70 μL of working solution (XTT reagent) were added directly to each well. Cells were incubated for 4h at 37°C in 5% of CO2 and 95% of humidity. After incubation, the absorbance was measured in the plate reader (Tecan) at 450 nm and 660 nm to eliminate the background signal contributed by cell debris or other non-specific absorbance. Negative controls with non-treated cells were included as blanks.

The percentage of cell viability was calculated using the following formula:

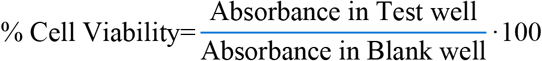

### 2.14 *C. acnes* EVs incubation in the outer skin layer model, the sebum model and the broken skin barrier model

*C. acnes* A1, H1 and H2 EVs were incubated with the immortalized HaCaT, SZ95 sebocytes and Jurkat cell lines (Fig. S2). For this purpose, 2·10^5^ cells/mL were seeded in a 12-well plate for the three cell lines. After 24h, all cell types were incubated for 24h with *C. acnes* A1, H1 and H2 EVs. The three plates were divided into three rows - one for each *C. acnes* phylotype - and each row had a control well and different concentrations of EVs, namely 12.5 μg/mL, 25 μg/mL and 50 μg/mL.

### 2.15 Quantitative Real Time-PCR analysis (RT-qPCR)

Total RNA was extracted from HaCaT, SZ95 sebocytes and Jurkat cells using the miRNeasy Mini Kit (Qiagen) following the manufacturer’s recommendations. Purity and RNA concentration were measured by the absorbance ratio at 260 and 280 nm on the NanoDrop One spectrophotometer (Thermo Scientific).

RNA was reverse transcribed using the cDNA Reverse Transcription Kit (Thermo Fisher Scientific) in a final volume of 20 μL following the manufacturer’s protocol. The retro transcription reaction was performed in the ProFlex PCR System (Applied Biosystems). RT-qPCR reactions were performed on the QuantStudio 7 Flex Real-Time PCR System (Applied Biosystems) using SYBR Green PCR Master Mix (Applied Biosystems) and cell-type specific primers. See the list of primers in Table S4 of the Supporting Information. A control reaction was set up with water in which no RNA was present. The 2^-ΔΔCt^ method was used to normalise expression results. The values of the housekeeping CREBBP gene were used to standardise the values obtained for each of the genes being studied.

### 2.16 Multiplex panel (Eve Technologies)

To validate the results obtained in the RT-qPCR, a Human High Sensitivity Plex Discovery Assay was performed (Eve Technologies). The targets of interest were selected from an extensive list of analytes and the company allowed multiplexing the previously selected targets together in a single assay. Among all the analytes, the ones that stood out for us in the present project were cytokines and chemokines.

### 2.17 Evaluation of sebum-reducing potential of *C. acnes* A1, H1 and H2 EVs in the acne-mimicking skin model

For the acneic model, human iPSC-derived sebocytes (PCi-SEB) were incubated for 5 days with maturation supplement. After that, the sebum production was induced by 48h of treatment with 5 μM of arachidonic acid (AA5). Then, cells were treated with 12.5 μg/mL of *C. acnes* A1, H1 and H2 EVs for 24h and cells were fixed until sebum production determination. In parallel, non-treated cells were used as negative control. For lipid droplets detection, cells were stained with the fluorescent marker BODIPY 493/503 (Sigma Aldrich) following manufacture’s recommendations. The nucleus was stained with DAPI. Samples were analysed by confocal fluorescence microscopy.

### 2.18 Statistical analyses

Statistical analyses were performed with GraphPad software. All tests were repeated independently at least three times in triplicate. The values for all measurements are presented as the mean ± standard deviation (SD). Differences between more than two groups were assessed using one-way or two-way ANOVA followed by Dunnett’s test. Data with a p value less than 0.05 and less than 0.001 were considered statistically significant and statistically highly significant, respectively.

## 3. RESULTS

### 3.1 Isolation and visualization of *C. acnes* EVs

*C. acnes* H1 EVs were isolated from cell-free culture supernatants (Fig. 1) and evaluated by negative stain-TEM. Images displayed spherical vesicles approximately 70-100 nm in diameter (Fig. 2A). To validate the TEM results and to obtain additional information, NTA which is a more precise imaging technique, was carried out in *C. acnes* A1, H1 and H2 EVs. This last analysis provided high-resolution particle size distribution profiles and also concentration measurements. Fig. 2B shows the size distribution graphs of the different *C. acnes* EVs. On the other side, values obtained after NTA revealed that H2 EVs have the largest mean vesicle size. However, in terms of concentration, A1 is the phylotype with the highest EV secretion yield rate (3.92 · 10^12^ EVs/mL). Moreover, recordings of the different types of *C. acnes* EVs were acquired from the NTA analysis (see Videos).

**Fig. 1.**
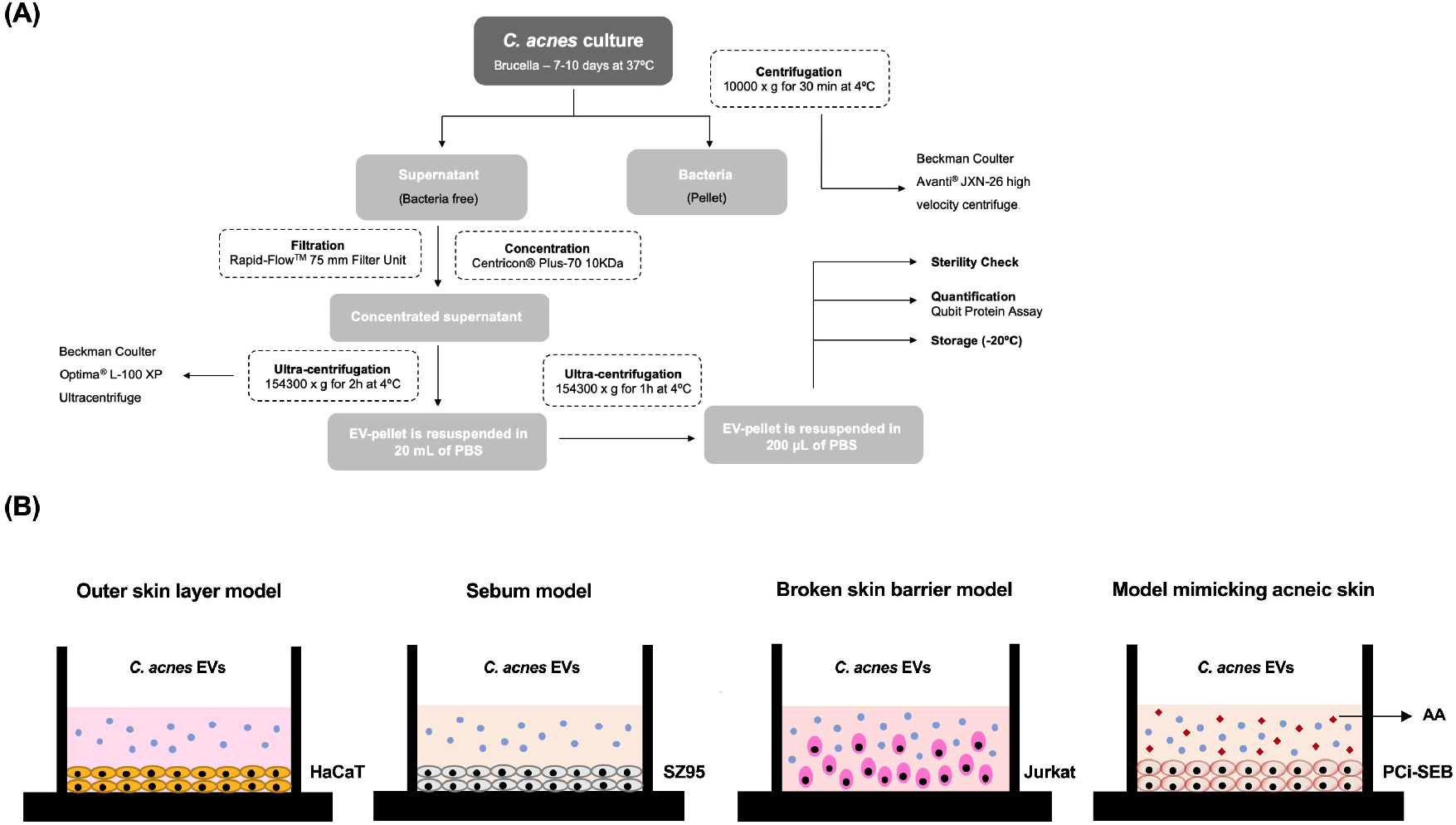
EVs isolation protocol and *in vitro* experimental design. (A) Process to isolate C. acnes-derived extracellular vesicles (EVs). (B) Schematic presentation of the different skin in vitro models: the outer skin layer model with HaCaT cell line, the sebum model with SZ95 sebocyte cell line, the broken skin barrier model with Jurkat cells and the acneic skin model with PCi-SEB cells using AA to induce sebum production.

**Fig. 2.**
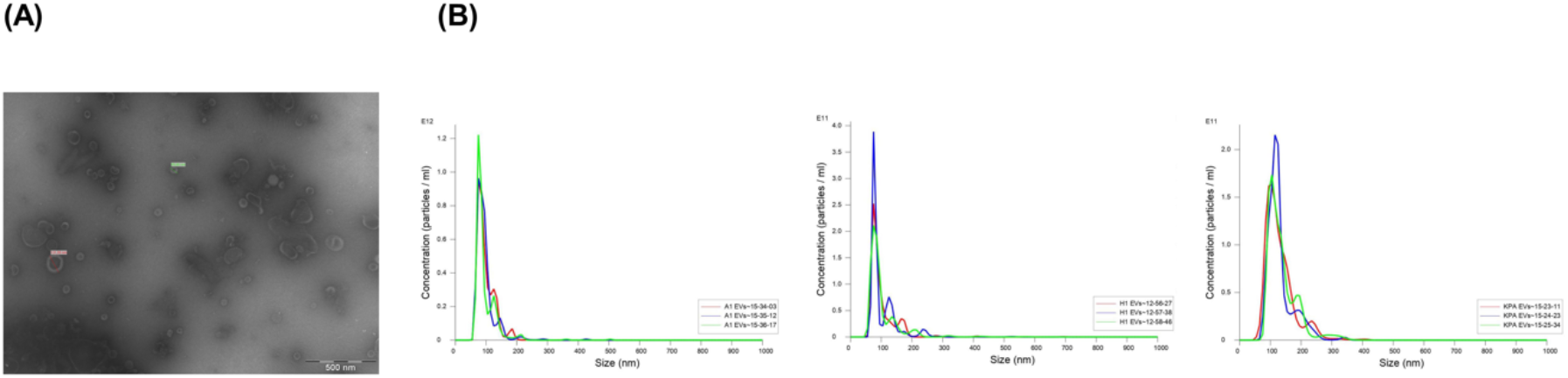
Visualization of *C. acnes* EVs by different microscopy techniques. (A) Transmission Electron Microscopy of C. acnes H1 EVs (bar 500 nm). (B) Nanoparticle Tracking Analysis of *C. acnes* A1, H1 and H2 EVs. Prior to the measurement, the samples were diluted in PBS buffer at 1:5000, 1:1000 and 1:1000 respectively. EVs have a mean diameter of 96.8 ± 0.5 nm, 100.7 ± 0.7 nm and 133.0 ± 2.3 nm respectively. EV-concentration is expressed as a number of particles per mL on the y axis. All data represent the mean of three independent experiments ± standard error.

### 3.2 Proteomic analysis comparison between *C. acnes* A1, H1 and H2 EVs

SDS-PAGE analysis showed that EV triplicates of each phylotype display similar protein profiles with slight differences (Fig. 3A), which is consistent with the suggested variability of protein cargo between batches (Gandham et al., 2020; Hartjes et al., 2019a, 2019b; Paganini et al., 2019). Furthermore, it should be noted that, although the same amount of protein was loaded into the gel for the EVs of each phylotype (7 μg), the diversity of proteins is much higher in A1 EVs (clade IA1) than in H1 and H2 EVs (clade IB). These findings are in line with the ones obtained by Chudzik et al.

**Fig. 3.**
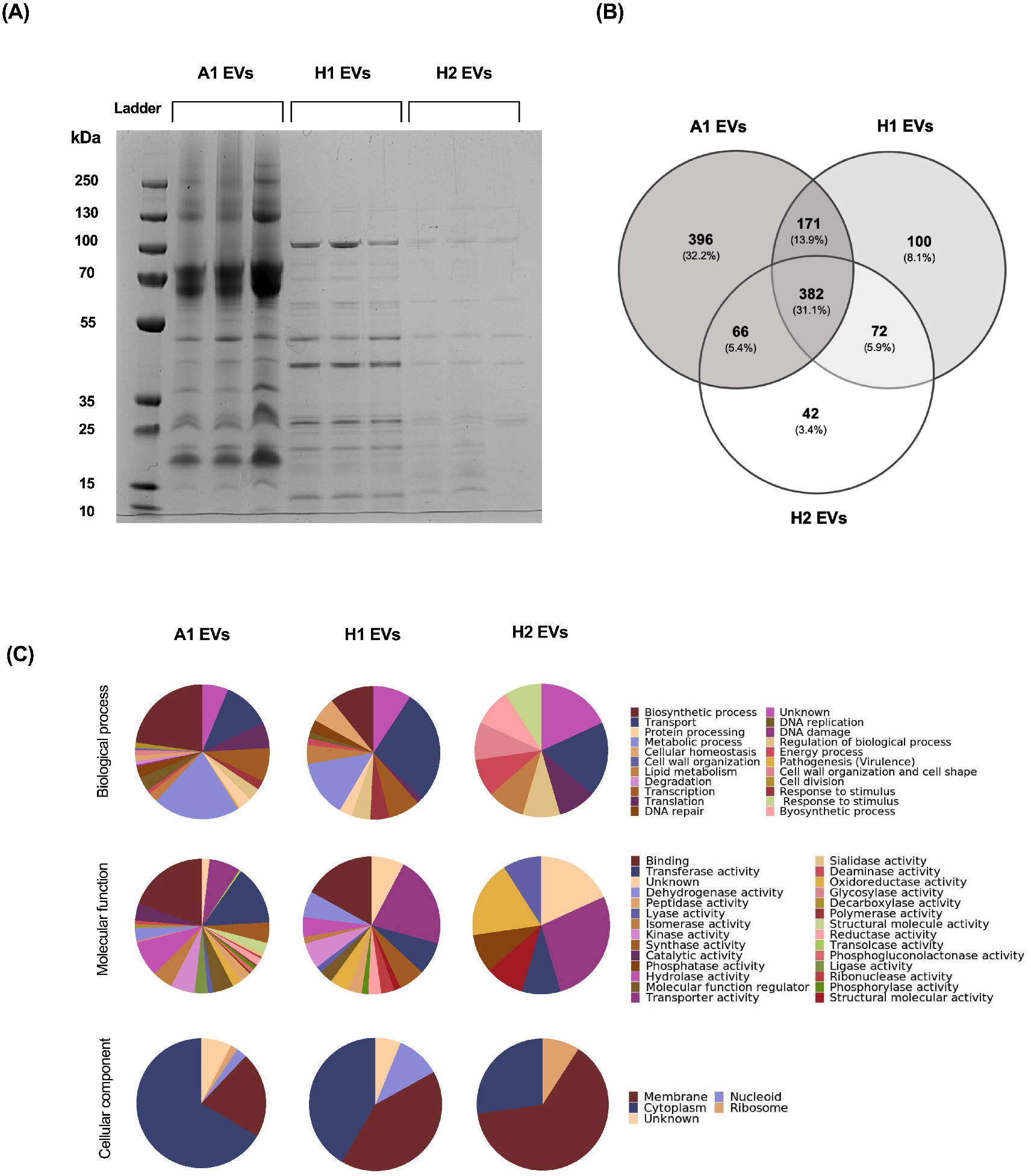
Proteomic characterization of *C. acnes* A1, H1 and H2 EVs. (A) Proteomic fingerprint of *C*. *acnes* EVs. SDS-PAGE gel showing protein bands of triplicates from EVs of different phylotypes. A concentration of 7 μg of total amount of protein was loaded in a NuPage 4 to 12% Bis-Tris gel and stained with Coomassie Brilliant Blue R-250 staining solution. (B) Venn diagram of the proteins contained in *C. acnes* A1, H1 and H2 EVs. The number of overlapping proteins between the different phylotypes is indicated. All data is obtained from three independent experiments. (C) Gene Ontology analysis of *C. acnes* A1, H1 and H2 EVs preparations. Biological process, Molecular function and Cellular components of the identified vesicular proteins in *C. acnes* EVs are presented here. For each *C. acnes* phylotype, three independent batches were analysed in the proteomic analysis.

To characterize the protein profile of *C. acnes* A1, H1 and H2 EVs, the protein content was isolated and examined by mass spectrometry by the UPF/CRG Proteomics Facility. As Fig. 3B illustrates in a Venn diagram, the three phylotypes share 382 proteins. Additionally, a pairwise comparison was performed, and the results showed that A1 and H1 share 171 proteins, A1 and H2 share 66 proteins and H1 and H2 share 72 proteins.

Finally, observing the proteins exclusive to each phylotype it is shown that A1 EVs have the highest number of high-confidence proteins identified, namely 396, followed by H1 and H2 EVs. Using the UniProt database, a Gene Ontology (GO) and bioinformatics analysis were conducted (Fig. 3C). In terms of biological processes, *C. acnes* A1 EVs were significantly enriched in proteins participating in biosynthetic processes (22.77%) and metabolic processes (20.98%). Regarding the molecular functions, binding (19.64%), transferase activity (14.29%) and hydrolase activity (8.48%) were the most prevalent. For biological processes, proteins linked to *C. acnes* H1 EVs were significantly enriched in transport (27.69%) and metabolic processes (13.85%). Regarding the molecular functions, transporter activity (21.54%) and binding (16.92%) were noteworthy. The predominant proteins involved in the biological processes of *C. acnes* H2 EVs were linked to transport (18.18%) and response to stimulus (9.09%). Regarding the molecular functions of the vesicular proteins found in this phylotype, transporter activity (27.27%), and oxidoreductase activity (18.18%) were the most prevalent. For all three types of EVs, in terms of cellular components, the highest percentage of proteins were membrane-associated proteins, followed by cytoplasmic proteins. Also, approximately 6-18% of the total number of proteins were not properly classified as they were unknown or poorly characterized. The identified proteins in *C. acnes* A1, H1 and H1 EVs are listed respectively in Tables S1, S2 and S3 of the Supporting Information.

### 3.3 EVs from *C. acnes* A1 phylotype are internalized into keratinocytes and sebocytes

*C. acnes* EVs internalization in HaCaT keratinocytes and SZ95 sebocytes was confirmed by confocal fluorescence microscopy at 24h of incubation with rhodamine B-R18-labeled A1 EVs (2 μg/ well). Membranes were labelled with WGA and nuclei with Hoechst. As anticipated, no red signal was observed in untreated control cells (Fig. 4).

**Fig. 4.**
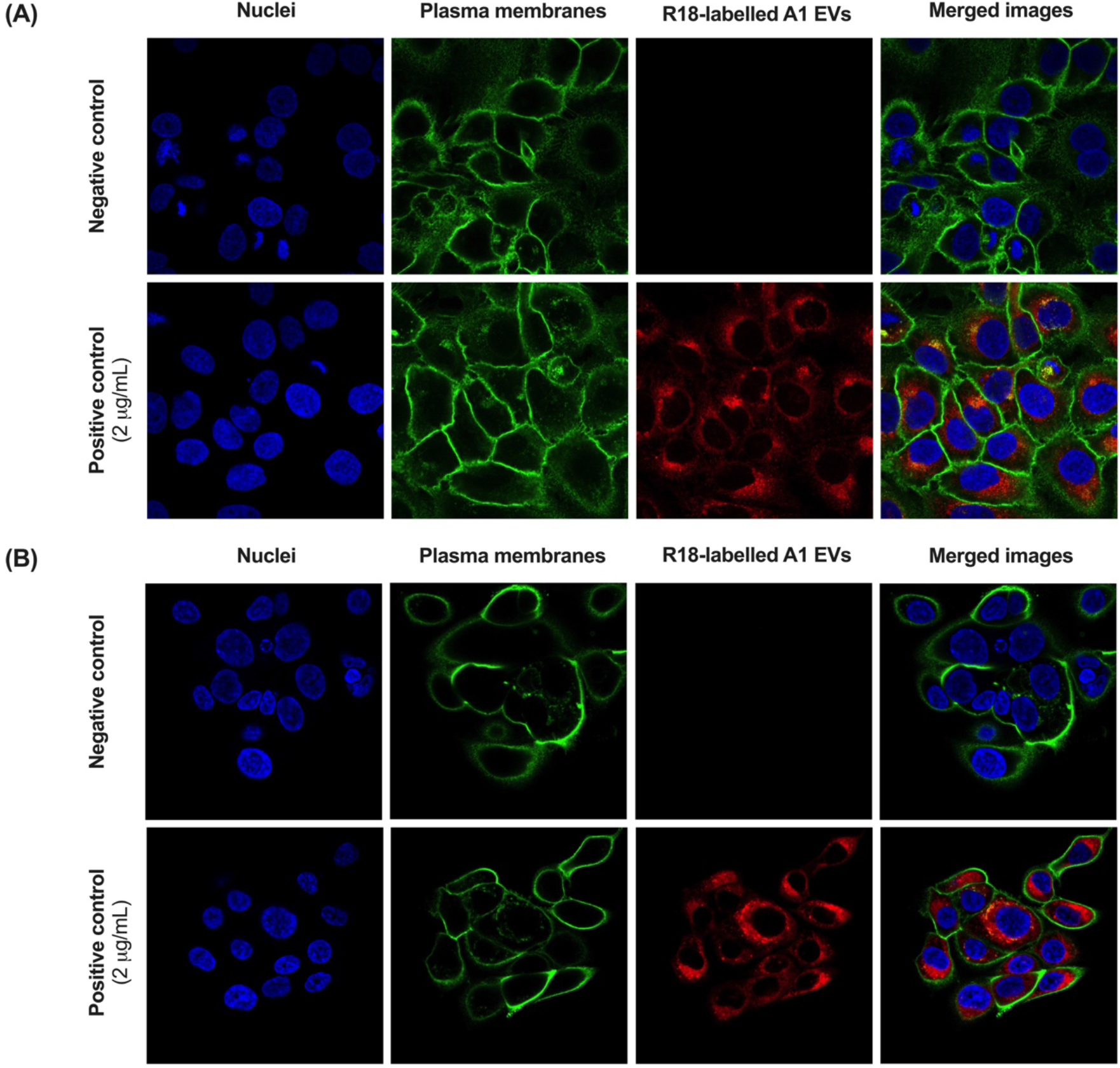
Uptake of *C. acnes* A1 EVs in (A) HaCaT cell line and (B) SZ95 cell line. Visualization of internalized EVs by confocal fluorescence microscopy. Both cell lines were incubated for 24h at 37°C with rhodamine B-R18-labelled A1 EVs (2 μg/ well).

### 3.4 From a concentration of 50 μg/mL and below, *C. acnes* EVs do not appear to affect cell viability in the two different skin models tested

A cytotoxicity assay was performed in HaCaT and SZ95 cells. In both tests, *C. acnes* A1, H1 and H2 EVs were added at different doses (1.56 μg/mL, 3.125 μg/mL, 6.25 μg/mL, 12.5 μg/mL, 25 μg/mL, 50 μg/mL, 100 μg/mL and 200 μg/mL). As is shown in Fig. 5A, in the HaCaT model cell viability is not reduced in any condition.

**Fig. 5.**
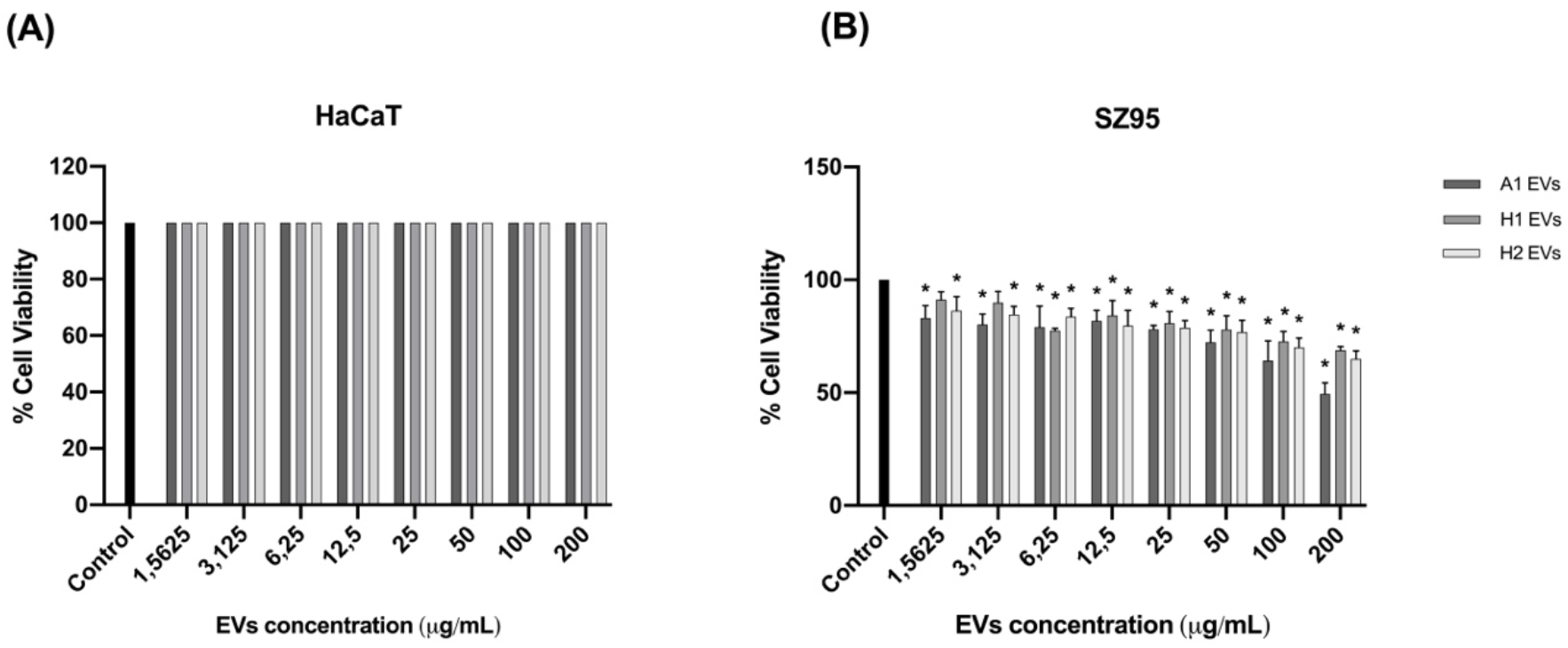
Cytotoxicity assay of (A) HaCaT cell line and (B) SZ95 sebocyte cell line treated with different doses of *C. acnes* A1, H1 and H2 EVs. All data are presented as mean ± standard deviation (SD) of triplicate measurements (*p ≤ 0.05 versus non-stimulated controls).

On the other hand, in the SZ95 model cell viability is reduced to 50-70% when concentrations of 200 μg/mL and 100 μg/mL of EVs are added (Fig. 5B). Nonetheless, in this model when adding concentrations of 50 μg/mL and below, viability is lowered by approximately 15-20% in A1 and H2 EVs but is maintained nearly at 100% in H1 EVs.

### 3.5 EVs released by *C. acnes* A1, H1 and H2 phylotypes modulate differently the expression of biomarkers related to the immune system, oxidative stress, skin barrier and sebum production

Different *in vitro* skin models were used to evaluate the immunomodulatory properties of direct stimulation with a range of concentrations of *C. acnes* A1, H1 and H2 EVs (Fig. 1B). In this sense, HaCaT, SZ95 and Jurkat immortalized cell lines were used (Colombo et al., 2017; Hewitt et al., 2013; Kühbacher et al., 2017; Luongo et al., 2014; Nikolakis et al., 2015; Sutterby et al., 2022; Törőcsik et al., 2021). In all *in vitro* models, cells were stimulated with *C. acnes* EVs for 24h, and after that, the expression level of different regulatory genes (immune system, skin barrier and oxidative stress) was measured by RT-qPCR. As Fig. 6A shows, A1 EVs trigger a higher stimulation of the pro-inflammatory cytokines Interleukin-8 (IL-8) and TGFβ-1, and the prostaglandin COX-2 which is linked to oxidative stress, compared to H1 or H2 EVs. Regarding genes involved in the reinforcement of the skin barrier, such as occludin, all concentrations of H2 EVs and high concentrations of H1 EVs (25 μg/mL and 50 μg/mL) were the only ones able to induce a significant increase in the mRNA expression compared to the negative control. However, for claudin-1, all the conditions seem to activate the expression of this gene, except low doses of H2 EVs, compared to untreated cells. In contrast, expression levels of MMP-2, a matrix metalloproteinase implicated in tissue remodelling (Manuel & Gawronska-Kozak, 2006), did not show marked differences for any of the EVs treatment with respect to the control. Regarding the skin immune system model, as Fig. 6B shows, only A1 EVs triggered a significant expression of the pro-inflammatory cytokine IL-8 compared to the control, while only H1 EVs and high doses of H2 EVs (50 μg/mL) had substantial anti-inflammatory cytokine IL-10 expression. However, for the pro-inflammatory TNFα expression, none of the EVs conditions tested were able to modulate its mRNA levels. As illustrated in Fig. 6C, A1 EVs are constantly triggering greater activation of the inflammatory cytokine IL-8. Otherwise, for the sebum regulator gene PLIN-2 none of the EVs tested were able to substantially stimulate the up-regulation of this gene expression compared to the untreated control.

**Fig. 6.**
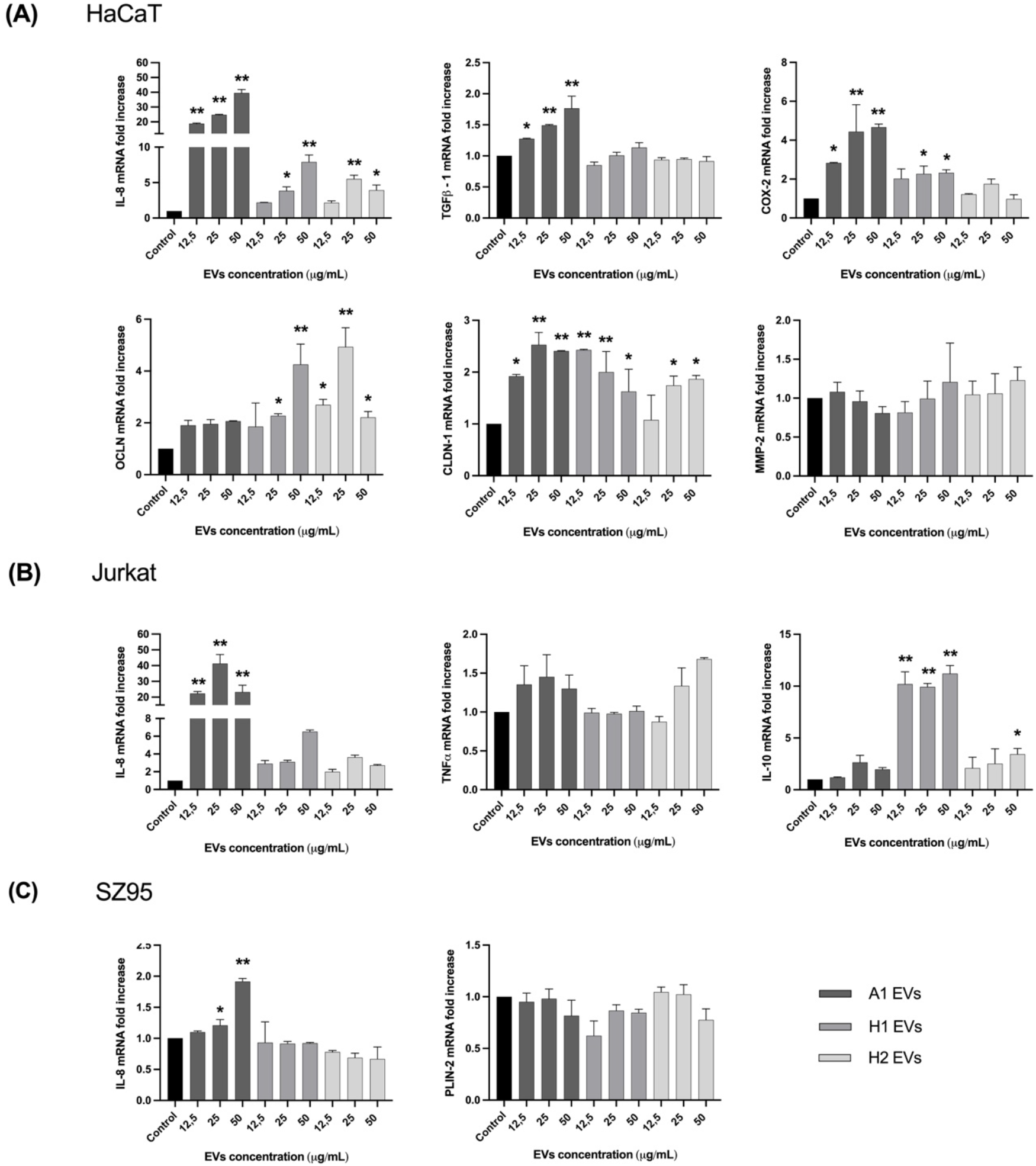
*In vitro* analysis with direct incubation of *C. acnes* EVs. Human cells were treated for 24h with different concentrations (12.5, 25 and 50 μg/mL) of *C. acnes* A1, H1 and H2 EVs. Total RNA was extracted and different biomarkers were assessed by RT-qPCR. (A) HaCaT (B) SZ95 and (C) Jurkat immortalized cell lines. All data are presented as mean ± standard deviation (SD) of triplicate measurements (*p ≤ 0.05, **p ≤ 0.001 versus non-stimulated controls).

### 3.6 EVs derived from *C. acnes* A1 phylotype induce a higher secretion of immunomodulatory mediators in three *in vitro* skin models compared to EVs from *C. acnes* H1 and H2 phylotypes

To confirm the immunomodulatory effects assayed by RT-qPCR in HaCaT, Jurkat and SZ95 *in vitro* models, after 24h of incubation with different concentrations of A1, H1 and H2 EVs, the protein secretion levels of some cytokines released in the SN were evaluated by Human High Sensitivity Plex Discovery Assay (Eve Technologies).

As shown in Fig. 7A in the HaCaT *in vitro* model, and following the same pattern as for IL-8 expression level, A1 EVs displayed the highest secretion of IL-8, IL-6, TNFα and GM-CSF compared to the rest of the EVs, especially when high concentrations of EVs were applied to the keratinocytes. Only 25 and 50 μg/mL for H1 EVs and 50 μg/mL for H2 EVs were able to induce a significant difference in IL-6 secretion compared to the negative control. In the immune system model, composed of Jurkat cells (Fig. 7B), high doses of A1 EVs were observed to trigger increased secretion of IL-8, IL-6 and TNFα compared to the other conditions, but in the case of GM-CSF, H1 and H2 EVs were the ones involved in the activation of this factor. Regarding the SZ95 model (Fig. 7C), IL-8 stimulation correlates with the gene expression data obtained previously. Overall, it can be observed that A1 EVs consistently trigger significant activation of the inflammation-related factors IL-8, IL-6, TNFα and GM-CSF, compared to the negative control. In the case of H1 EVs, only 25 and 50 μg/mL were able to induce significant IL-8 release compared to control.

**Fig. 7.**
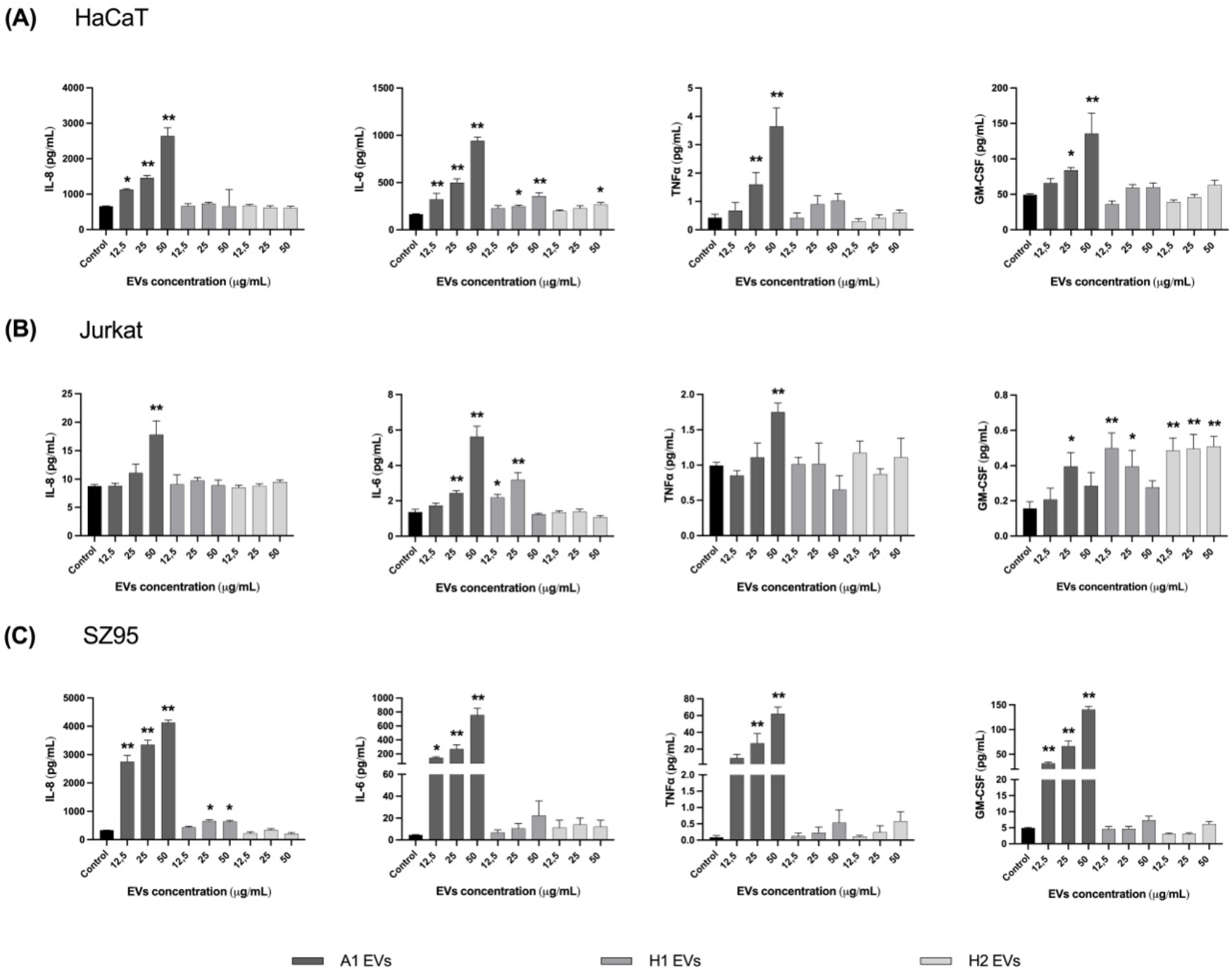
Total protein concentration secreted into the SN after direct incubation with *C. acnes* EVs. Human cells were treated for 24h with different concentrations (12.5, 25 and 50 μg/mL of *C. acnes* A1, H1 and H2 EVs. Filtered SN was evaluated by Multiplex analysis to measure protein secreted in it. Ranges for the standard curve for each of the biomarkers were: GM-CSF: 0.31 - 4978.62 pg/mL, TNF alpha: 0.11 - 1699.58 pg/mL and IL-6: 0.05 - 768.83 pg/mL. (A) HaCaT (B) SZ95 and (C) Jurkat immortalized cell lines were used. All data are presented as mean ± standard deviation (SD) of triplicate measurements (*p ≤ 0.05, **p ≤ 0.001 versus non-stimulated controls).

### 3.7 *C. acnes* H1 EVs showed to have a better skin sebum inhibition mechanism compared to A1 and H2 EVs

The acneic *in vitro* model consisted of a 48-hour treatment with AA5 as a sebum inducer followed by a 24-hour treatment with 12.5 μg/mL of *C. acnes* A1, H1 and H2 EVs. After AA5 incubation, a strong and significant increase in lipid level by about 263-fold was observed, compared to the negative control condition. When PCi-SEB were treated for 24h with 12.5 μg/mL of *C. acnes* A1, H1 and H2 EVs, they showed a 2.8, 4.3 and 3.6-fold reduction *versus* the AA5 condition, respectively. However, although results showed that the three types of EVs had a significant inhibitory effect on sebum production compared to the positive control, the best sebum inhibitory effect was obtained after H1 EVs treatment, with about 4-fold inhibition compared to AA5 condition (Fig. 8).

**Fig. 8.**
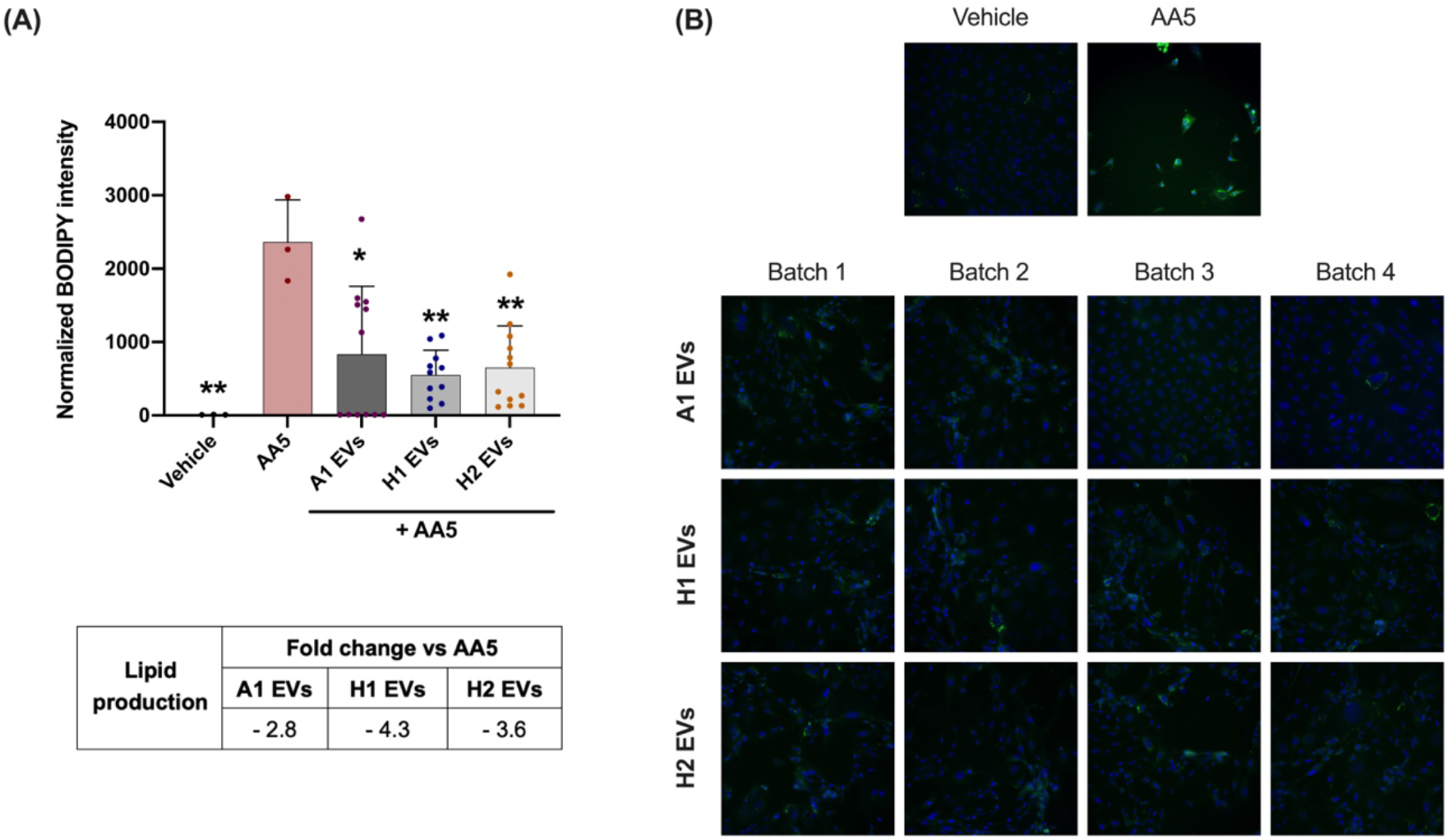
Reduction of lipid production was assessed after treating PCi-SEB with AA5 for 24h. PCi-SEB were treated during 48h with AA at 5 μM (AA5) to induce lipid production. Afterwards PCi-SEB were treated for 24h with 12.5 μg/mL of *C. acnes* A1, H1 and H2 EVs (A) Metadata analysis inter-EVs batches and (B) Fluorescence microscopy image of PCi-SEB stained with BODIPY 493/503; negative control (Vehicle), positive control (AA5). All data are presented as mean ± standard deviation (SD) of triplicate measurements (*p ≤ 0.05, **p ≤ 0.001 versus non-stimulated controls).

## 4. DISCUSSION

The Gram-positive bacterium *C. acnes* is one of the main representative genera presents in the human skin microbiota with a key and pivotal role in maintaining a healthy skin barrier. In the same way that occurs in the intestine, communication between C. acnes and host cells it is proposed to be *established* via soluble mediators and EVs that can diffuse through the different layers of the skin (epidermis, dermis and hypodermis). Specifically, EVs act as nano-carriers of proteins, metabolites and nucleic acids upon interaction and internalization into host cells. Therefore, it has been suggested that, considering the cargo of *C. acnes* EVs, they will have a different role, being responsible on the different effects on skin alterations. *C. acnes* strains are currently classified into eight phylotypes (IA1, IA2, IB1, IB2, IB3, IC, II and III), being the IA1 and IA2 clades preferentially found in acne lesions whereas phylotype IB is linked to a healthy skin without sign of acne. In this context, where acne vulgaris is the 8th most prevalent disease, with a 9.4% world population affection and several partially effective therapeutic compounds and procedures in the market (Tuchayi et al., 2015), the objective of this project has been focused on studying the role of EVs secreted by A1 (IA1), H1 (IB) and H2 (IB) *C. acnes* phylotypes as a safety biotherapy in cutaneous disorders such acne vulgaris.

EV research remains a major challenge due to their intrinsically complex biogenesis, their great heterogeneity in terms of size and the natural variations between batches encountered during their production (Gandham et al., 2020; Hartjes et al., 2019; Paganini et al., 2019). Nevertheless, the nature (small, non-synthetic and non-replicative) of these structures makes them unique to overcome important barriers found in the current dermatology and cosmetic field as it is the skin penetrance, side effects derived from chemicals and maintaining the skin microbiota symbiosis. However, despite this good prognose of EVs used as bio-carriers, what will determine a successful, harmful or poor functionality in terms of acne vulgaris pathology, depends on the specific cargo of the EVs, which is directly related to the bacterium from which they derive. Therefore, as the main components are proteins, a proteomic characterization of *C. acnes* EVs was performed to determine the relationship of their protein cargo with the skin microbiota. A Venn diagram identified, in agreement with Chudzik et al., 2022, that A1 EVs contain the highest number of proteins, in our case almost four times more proteins than in H1 EVs and nine times more than in H2 EVs. These results were validated with an SDS-PAGE, where the band profile was different between phylotypes (Article). This could be associated with the extra number of pathogenic proteins that EVs of A1 phylotype harbour in contrast to H1 and H2 EVs. In fact, McDowell A. et al, 2013 found that certain strains of *C. acnes* (specifically type IA1) were capable of secreting several virulence factors that enhanced the inflammatory response. However, there are no genetic studies of the *C. acnes* strains used in this work, but as it has been showed previously, H1 has a protective effect against acne vulgaris development. Thus we could assume that these two strains could have lost the virulence factors during evolution. Following this assumption, 25 virulence factors (related to adhesion, biofilm formation, etc.) were found during the mass spectrometry analysis for A1 EVs while only 2 were attributed to H1 and 0 for H2 *C. acnes* EVs.

Regarding shared proteins by all the *C. acnes* EVs types, the proteomic analysis revealed that a large amount of proteins related to facilitate the transfer of multiple proteins involved in biochemical processes and metabolism between cells, were present in EVs, validating the carrier function of these structures. Moreover, EVs are used as a way of exchange cell surface substances and improve bacterial survival during infections (Chen et al., 2022), and in this sense we could identify several proteins used in antibiotic resistance. These findings suggest that *C*. acnes EVs are also involved in bacterial survival against external threats. In addition, due to the nature and sebum oily niche of *C. acnes*, common proteins involved in lipolysis such as lysophospholipase and triacylglycerol lipase, were identified in the EVs. Although these enzymes are generally considered putative virulence factors, lipases are necessary for *C. acnes* to maintain healthy skin by metabolizing sebum and releasing fatty acids, which are necessary to keep a proper skin acidity that acts as a natural barrier against harmful pathogens, providing innate skin immunity (Kim et al., 2020).

Considering the analysis of *C. acnes* phylotype-exclusive proteins for A1, H1 and H2 EVs, important differences were found in accordance with previous studies, pointing to *C. acnes* phylogroup A1 (IA1) as an acne-associated group and *C. acnes* phylogroup H1 (IB) as a beneficial and skincare group (Agak et al., 2018; Choi et al., 2018b; Chudzik et al., 2022; Fitz-Gibbon et al., 2013; Liu et al., 2014; Lomholt & Kilian, 2010; McLaughlin et al., 2019; Nakase et al., 2021; O’Neill & Gallo, 2018; Paetzold et al., 2019). As for A1 EVs, its protein profile revealed a great number of proteins involved in bacterial competition and antibiotic resistance (hydrolase, Ppx/GppA phosphatase family, Metallo-ß-lactamase domain protein and histidine kinase) (McLaughlin et al., 2019), biofilm formation (glycosyltransferase and serine/threonine-protein kinase) and virulence (porphyrin, sialidase, HtaA domain protein, phosphoesterase, hydrolase and sigma factor SigA) which were not detected in H1 or H2 EVs. Regarding the exclusive proteins of H1 EVs, different proteins directly involved in fatty acid catabolism were found, such as acyl-CoA dehydrogenase (Swigoňová et al., 2009) and a putative two-component sensor kinase which upregulates the hydrolysis of sebum triglycerides secreting free fatty acids. Indirectly, the N-acetyl-gamma-glutamyl-phosphate reductase, a protein involved in the L-Arginine biosynthetic pathway was also found as part of H1 EVs. In this regard, L-arginine helps to protect the skin from free radicals, increases the skin’s visible hydration levels, and potentially supports collagen production (Gad, 2010). All these results indicate that EVs and bacteria from which they derive, share functionalities, and it point to EVs as one of the mechanisms of action by which the specific *C. acnes* strains exert their effect, as it has been previously described by Choi et al., 2018b or Briaud & Carroll, 2020.

Then, the next step in our study was to show if the presence of unique peptides in the different types of *C. acnes* EVs could translate, as expected, in different functionalities in the host. For this purpose, an in vitro system representing the pathophysiology derived from acne was used. During acne vulgaris development, as Mattii et al., 2018 showed, not only the external skin layer is affected, but sebocytes also exhibit a pivotal role. Nonetheless, unlike most of the publications using skin models are based only in keratinocytes, in this work, other cell types were taken into consideration, as sebocytes and immune cells, to have a closer picture of what is happening in real skin during acne vulgaris. To do that, different in vitro assays were carried out. Regarding cytotoxicity assays, they revealed that doses of 50 μg/mL and below of *C. acnes* EVs did not appear to affect cell viability either in the HaCaT model or in the SZ95 sebocyte model, thus deeming them as safe to be tested in the in vitro skin models. Comparing both cell types, it was also observed that sebocytes were more responsive than keratinocytes. This can be explained because sebocytes inhabit in the sebaceous gland, which is located in the dermis, protected from external microbes in contrast to keratinocytes exposed and in constantly interaction with skin microorganisms.

Then, the different EVs were directly incubated for 24h in the different in vitro models: a) the outer layer skin model consisted of HaCaT cells to mimic the epidermis composed mainly of keratinocytes; b) the sebum model consisted of SZ95 cells to mimic the sebaceous gland environment and c) the broken barrier skin model, consisting of Jurkat cells, was chosen to get an idea of when the skin barrier is disrupted, one of the severe attributes of inflammatory skin diseases. The expression levels of different regulatory genes were measured by RT-qPCR and results displayed that, as previously was showed by Nagy et al., 2005, in all in vitro skin models, A1 EVs were systematically triggering increased activation of pro-inflammatory cytokines (Nagy et al., 2005). This could be due to the presence of a large number of virulence factors found in the proteomic analysis for A1 EVs. On the other side, in the broken skin barrier model, H1 EVs and 50 μg/mL doses of H2 EVs induced a significant expression of the anti-inflammatory cytokine IL-10, reinforcing the possible protective role of these two non-pathogenic strains during acne vulgaris development.

To cross-check the immunomodulatory effects assayed by gene expression, the protein secretion levels of some cytokines released in the SN of the *in vitro* models were evaluated and it was confirmed that EVs derived from *C. acnes* A1 induce a higher secretion of immunomodulatory mediators in the three *in vitro* skin models compared to *C. acnes* H1 and H2 EVs. According to the RT-qPCR results, it was determined that A1 EVs consistently triggered a higher activation of the inflammation-related factors IL-8, IL-6, TNFα and GM-CSF, compared to H1 and H2.

The pathophysiology of acne includes, apart from local inflammation, a hyperproduction of sebum by sebocytes which causes specific strains of *C. acnes* to overgrow causing a dysbiosis and triggering the development of acne (Paetzold et al., 2019; Rozas et al., 2021). In fact, one of the most prescribed treatments for dermatologists against acne is based on isotretionin, a retinoid which causes the apoptosis of skin sebocytes to reduce the amount of sebum (Zouboulis, 2006). However, the regulation of sebum during acne development modulated by the skin microbiota is a very new topic and there are few publications on the subject. Therefore, an acneic-mimicking skin model with PCi-SEB and AA5 as sebum inductor, was used to explore the effect of *C. acnes* EVs on sebum regulation. After 24-hour treatment with 12.5 μg/mL of *C. acnes* A1, H1 and H2 EVs, results showed that the three types of EVs had a significant inhibitory effect on sebum production compared to the positive control, highlighting that the best inhibition was obtained after H1 EVs treatment, with inhibition by about 4 folds compared to the positive control. These results were expected and in accordance with the proteomic analysis that described H1 EVs as being charged with exclusive proteins with lipidomic effect.

To our knowledge, this is the first study in which proteomic characterization of three different phylotypes of *C. acnes* has been performed, in addition to the testing of three different *C. acnes* phylotypes in different *in vitro* skin models. This work has proved that *C. acnes* EVs are more than lipid spheres that act as mechanisms of action of the derived bacteria. Specially H1 EVs has been highlighted as the best bioactive nanocarrier, counteracting acne symptomatology from two flanks, inhibiting inflammation and modulating sebum production.

## 5. CONCLUSION

The findings of this project have shown that *C. acnes* EVs are key intermediators in the communication between skin microbiota and the host, having an essential role during skin alterations such as acne vulgaris. In this sense, these results point to H1 EVs as a potential, efficient, safe and based-natural treatment to prevent and ameliorate symptomatology associated with inflammation and oily skin. However, further scientific experiments need to be done including assays with primary cell lines, *in vivo* animal models, checking EVs stability in a gel cream and performing clinical tests to extend the knowledge and to have the whole picture of how *C. acnes* EVs act on the skin.

## ABBREVIATIONS

*C. acnes*: *Cutibacterium acnes*
EVs: extracellular vesicles
PBS: phosphate-buffered saline
SN: supernatant
TEM: transmission electron microscopy
NTA: nanoparticle tracking analysis
SDS-PAGE: sodium dodecyl sulfate-polyacrylamide gel electrophoresis
PBS: phosphate-buffered saline
CPS: cycles per second
DDA: data-dependent acquisition
AGC: auto gain control
HCD: high-energy collision dissociation
FDR: false discovery rate
XIC: extracted ion current
ATCC: American Type Culture Collection
DMEM: Dulbecco’s Modified Eagle Medium
FBS: Fetal Bovine Serum
EGFr: recombinant human epidermal growth factor
CaCl_2_: calcium chloride
RPMI: Roswell Park Memorial Institute medium
RT: room temperature
WGA: wheat germ agglutinin
RT-qPCR: quantitative real time-PCR analysis
PCi-SEB: human iPSC-derived sebocytes
AA5: 5 μM of arachidonic acid
SD: standard deviation
GO: Gene Ontology
IL: interleukin
CITE: biofilm matrix dynamics

## ETHICS APPROVAL AND CONSENT TO PARTICIPATE

Not applicable

## CONSENT FOR PUBLICATION

All authors give their consent for publication of this work in Microbiome as a research article.

## AVAILABILITY OF DATA AND MATERIALS

The raw data from proteomic analysis are included as Supporting Tables S1, S2 and S3.

The original R scripts for plotting the extracellular vesicle data are available on Bitbucket.

(https://bitbucket.org/synbiolab/fabrega_vesicles/src/master/)

## AUTHOR’S CONTRIBUTIONS

M.P. participated in the experimental laboratory work, data curation, formal analysis, writing the main manuscript text and preparing all figures. J.M. did the bioinformatic analysis, prepared figure 3 and helped with the writing of the original draft. L.T. and N.K. participated in the experimental laboratory work, specially helping to set up the in vitro assays and bacterial cultures. J.M. participated in the experimental laboratory work, developing the in vitro acneic model and preparing figure 8. M.G. contributed supervising the project and helping to write the main manuscript text. M.F. helped with the experimental laboratory work, data curation, formal analysis, writing the manuscript, doing the funding acquisition/project administration, and supervising the project. C,C,Z, provided its proprietary imoortalized human sebaceous gland cell line SZ95 and revised the initial version of the manuscript.

## ACKNOWLEDGEMENT

We want to express our deepest gratitude to Dr. Bernhard Paetzold for support in culturing *C. acnes* strains. Also, Dr. Jordi García-Ojalvo for mentoring this project. We also want to thank the Office Naval Research (Award N62909-18-1-2155) for giving us the opportunity to study how to work with anaerobic skin bacteria.

