## Supporting Information for "New insights into the role of *Cutibacterium acnes*-derived extracellular vesicles in inflammatory skin disorders"

Universitat Pompeu Fabra Barcelona

### SUPPORTING INFORMATION

#### Table S1. List of proteins identified in *C. acnes*-derived A1 EVs under anaerobic conditions

| Gene name | Accession number | Protein name | GO - Biological process | GO - Molecular function | GO - Cellular component |
|---|---|---|---|---|---|
| pyrK PPA1001 cbac_05440 | WP_002517886.1 | Dihydroorotate dehydrogenase B | Biosynthetic process | Binding | Membrane |
| cbac_03745 | WP_073842171.1 | PTS sugar transporter | Transport | Transferase activity | Membrane |
| PPA1099 cbac_05960 | WP_002514993.1 | PAC2 family protein | Protein processing | Unknown | Cytoplasm |
| pyrD cbac_05445 | WP_002517834.1 | Dihydroorotate dehydrogenase | Biosynthetic process | Dehydrogenase activity | Cytoplasm |
| PPA2168 cbac_11750 | WP_002516248.1 | Formimidoylglutamate deiminase | Metabolic process | Peptidase activity | Cytoplasm |
| cbac_02600 | WP_002531152.1 | Copper homeostasis protein CutC | Cellular homeostasis | Binding | Unknown |
| folP PPA0271 cbac_01520 | WP_002517047.1 | Dihydropteroate synthase | Biosynthetic process | Binding | Cytoplasm |
| PPA0313 cbac_01745 | WP_002517195.1 | NAD-dependent malic enzyme | Metabolic process | Dehydrogenase activity | Cytoplasm |
| hisF HMPREF0675_4424 | WP_002516482.1 | Imidazole glycerol phosphate synthase subunit HisF | Biosynthetic process | Lyase activity | Cytoplasm |
| PPA0464 cbac_02555 | WP_002530084.1 | Sugar phosphate isomerase/epimerase | Biosynthetic process | Isomerase activity | Membrane |
| PPA0464 cbac_02555 | WP_002530084.1 | Sugar phosphate isomerase/epimerase | Biosynthetic process | Isomerase activity | Cytoplasm |
| PPA1005 cbac_05460 | WP_002515547.1 | Fructosamine kinase | Protein processing | Kinase activity | Cytoplasm |
| cbac_02675 | WP_002518785.1 | Phosphoserine transaminase | Biosynthetic process | Transferase activity | Cytoplasm |
| ddl cbac_12720 | WP_002515704.1 | D-alanine--D-alanine ligase | Cell wall organization | Binding | Cytoplasm |
| thiE cbac_04840 | WP_002531442.1 | Thiamine-phosphate synthase | Biosynthetic process | Binding | Cytoplasm |
| PPA0303 cbac_01695 | WP_002531197.1 | Uroporphyrinogen-III synthase | Biosynthetic process | Synthase activity | Cytoplasm |
| HMPREF0675_4140 | WP_002515011.1 | Histidine triad domain protein | Transport | Catalytic activity | Membrane |
| cobM PPA0421 cbac_02310 | WP_002515099.1 | Precorrin-4 C | Biosynthetic process | Transferase activity | Unknown |
| hutI HMPREF0675_5232 | WP_002516336.1 | Imidazolonepropionate | Metabolic process | Binding | Cytoplasm |
| PPA1225 cbac_06630 | WP_002524933.1 | Pyridoxal kinase | Biosynthetic process | Kinase activity | Cytoplasm |
| PPA1015 cbac_05515 | WP_002515534.1 | Aldose epimerase | Metabolic process | Binding | Membrane |
| PPA1982 cbac_10760 | WP_002530665.1 | Phosphatase PAP2 family protein | Lipid metabolism | Phosphatase activity | Membrane |
| PPA0525 cbac_02915 | WP_002516665.1 | DNase | Degradation | Hydrolase activity | Cytoplasm |
| PPA1009 cbac_05480 | WP_002513709.1 | Possible transcriptional regulator | Transcription | Molecular function regulator | Cytoplasm |
| PPA1629 cbac_08845 | WP_002517032.1 | 6-phosphogluconate dehydrogenase, decarboxylating | Metabolic process | Dehydrogenase activity | Cytoplasm |
| PPA2286 cbac_12435 | WP_002515771.1 | Phosphoglucomutase | Metabolic process | Transferase activity | Cytoplasm |
| PPA0106 cbac_00580 | WP_002512819.1 | ABC transporter substrate-binding protein | Transport | Binding | Membrane |
| tsaB PPA1784 cbac_09710 | WP_002531267.1 | Peptidase, family M22 | Translation | Transferase activity | Membrane |
| tsaB PPA1784 cbac_09710 | WP_002531267.1 | Peptidase, family M22 | Translation | Transferase activity | Cytoplasm |
| idi cbac_11445 | WP_002530756.1 | Isopentenyl-diphosphate Delta-isomerase | Biosynthetic process | Hydrolase activity | Cytoplasm |
| PPA2156 cbac_11695 | WP_002530776.1 | Dipeptide ABC transporter ATP-binding protein | Transport | Transporter activity | Membrane |
| murQ HMPREF9578_00165 | WP_002513573.1 | N-acetylmuramic acid 6-phosphate etherase | Metabolic process | Lyase activity | Cytoplasm |
| PPA2214 cbac_12030 | WP_002530902.1 | Electron transfer flavoprotein subunit alpha | Metabolic process | Transferase activity | Unknown |
| HMPREF9206_1221 | WP_002512973.1 | Phosphoglycerate mutase family protein | Metabolic process | Catalytic activity | Cytoplasm |

| | | | | | |
|---|---|---|---|---|---|
| menC PPA0902 cbac_04915 | WP_002515608.1 | o-succinylbenzoate synthase | Biosynthetic process | Synthase activity | Cytoplasm |
| cmk HMPREF0675_4273 | WP_002516760.1 | Cytidylate kinase | Metabolic process | Kinase activity | Cytoplasm |
| metK HMPREF0675_4255 | WP_002517758.1 | S-adenosylmethionine synthase | Metabolic process | Binding | Cytoplasm |
| cbac_03780 | WP_002530937.1 | Exo-alpha-sialidase | Lipid metabolism | Sialidase activity | Cytoplasm |
| PPA1781 cbac_09695 | WP_002514786.1 | 8-oxo-dGTP diphosphatase | DNA repair | Hydrolase activity | Cytoplasm |
| PPA1544 cbac_08415 | WP_002514359.1 | Conserved protein | Protein processing | Binding | Cytoplasm |
| PPA1950 cbac_10560 | WP_002523392.1 | Cell division protein DedD | Metabolic process | Deaminase activity | Membrane |
| coaD HMPREF9578_02029 | WP_002514272.1 | Phosphopantetheine adenylyltransferase | Biosynthetic process | Transferase activity | Cytoplasm |
| PPA2116 cbac_11450 | WP_002530757.1 | Serine protease | Biosynthetic process | Peptidase activity | Membrane |
| PPA0443 cbac_02415 | WP_002516675.1 | Cobalamin-binding protein | Transport | Binding | Cytoplasm |
| HMPREF0675_3579 | WP_002517279.1 | Transcriptional regulator, TetR family | Transcription | Molecular function regulator | Nucleoid |
| HMPREF0675_5283 | WP_002516338.1 | FAD dependent oxidoreductase | Metabolic process | Oxidoreductase activity | Cytoplasm |
| PPA1366 cbac_07465 | WP_002516499.1 | FAA hydrolase family protein | Metabolic process | Catalytic activity | Cytoplasm |
| manA PPA0008 cbac_00045 | WP_002515893.1 | Mannose-6-phosphate isomerase | Metabolic process | Isomerase activity | Cytoplasm |
| cbac_06585 | WP_002516778.1 | HAD family hydrolase | Protein processing | Hydrolase activity | Cytoplasm |
| PPA1377 cbac_07530 | WP_002531311.1 | RNA helicase | Unknown | Binding | Cytoplasm |
| PPA0603 cbac_03350 | WP_007400924.1 | Putative transcriptional regulator | Transcription | Molecular function regulator | Unknown |
| HMPREF0675_5327 | WP_002518141.1 | Transcriptional regulator, MarR family | Transcription | Molecular function regulator | Cytoplasm |
| gyrA PPA0010 cbac_00055 | WP_002530866.1 | DNA gyrase subunit A | DNA replication | Binding | Cytoplasm |
| cbac_02560 | WP_002524726.1 | Sugar phosphate isomerase/epimerase | Metabolic process | Isomerase activity | Cytoplasm |
| cbac_04330 | WP_073885323.1 | HtaA domain protein | Translation | Binding | Membrane |
| PPA1181 cbac_06395 | WP_002516773.1 | Shikimate 5-dehydrogenase | Metabolic process | Dehydrogenase activity | Cytoplasm |
| prcB PPA1206 cbac_06530 | WP_002516776.1 | 20S proteasome beta-subunit | Metabolic process | Peptidase activity | Cytoplasm |
| PPA0947 cbac_05145 | WP_002513775.1 | DNA-binding response regulator | Transcription | Molecular function regulator | Cytoplasm |
| HMPREF0675_5322 | WP_002516444.1 | Putative 3-methyladenine DNA glycosylase | DNA replication | Binding | Nucleoid |
| HMPREF0675_5322 | WP_002516444.1 | Putative 3-methyladenine DNA glycosylase | DNA replication | Binding | Cytoplasm |
| PPA1371 cbac_07495 | WP_002516506.1 | Putative transferase | Biosynthetic process | Transferase activity | Unknown |
| PPA2273 cbac_12370 | WP_002519568.1 | Adenosine deaminase | DNA damage | Deaminase activity | Cytoplasm |
| gatA cbac_06100 | WP_002531385.1 | Glutamyl-tRNA | Translation | Hydrolase activity | Cytoplasm |
| PPA1631 cbac_08855 | WP_002517011.1 | Thioesterase | Regulation of biological process | Catalytic activity | Cytoplasm |
| cbac_10785 | WP_002530669.1 | ATPase | Energy process | Hydrolase activity | Membrane |
| PPA1290 cbac_07010 | WP_002531340.1 | 2,3-diaminopropionate biosynthesis protein SbnA | Biosynthetic process | Synthase activity | Cytoplasm |
| fmt HMPREF0675_4257 | WP_002517988.1 | Methionyl-tRNA formyltransferase | Translation | Transferase activity | Cytoplasm |
| tmk HMPREF0675_3289 | WP_002517149.1 | Thymidylate kinase | Biosynthetic process | Kinase activity | Cytoplasm |
| ung PPA0558 cbac_03120 | WP_002519227.1 | Uracil-DNA glycosylase | DNA repair | Glycosylase activity | Cytoplasm |
| PPA1277 cbac_06910 | WP_002531349.1 | ABC transporter substrate-binding protein | Transport | Binding | Membrane |
| cbac_09495 | WP_041444224.1 | Phosphoesterase | Pathogenesis (Virulence) | Hydrolase activity | Cytoplasm |
| lysA PPA1259 cbac_06800 | WP_002516817.1 | Diaminopimelate decarboxylase | Biosynthetic process | Decarboxylase activity | Cytoplasm |
| rpoZ HMPREF1162_0201 | WP_002516821.1 | DNA-directed RNA polymerase subunit omega | Transcription | Polymerase activity | Nucleoid |
| HMPREF9578_00185 | WP_002516610.1 | ACT domain-containing protein | Metabolic process | Structural molecule activity | Cytoplasm |
| PPA0563 cbac_03150 | WP_011183699.1 | Conserved protein containing thioredoxin domain | Metabolic process | Reductase activity | Cytoplasm |

| | | | | | |
|---|---|---|---|---|---|
| cbac_01645 | WP_002523010.1 | PTS glucose transporter subunit IIB | Transport | Kinase activity | Membrane |
| PPA0548 cbac_03045 | WP_002516725.1 | Hydrolase, Ppx/GppA phosphatase family | Metabolic process | Hydrolase activity | Cytoplasm |
| PPA2167 cbac_11745 | WP_002516352.1 | Allantoate amidohydrolase (AAH, allantoinase) | Metabolic process | Hydrolase activity | Cytoplasm |
| PPA0814 cbac_04460 | WP_002519094.1 | Putative gluconeogenesis factor | Cell wall organization and cell shape | Transferase activity | Cytoplasm |
| map PPA1833 cbac_09965 | WP_002523458.1 | Methionine aminopeptidase | Protein processing | Peptidase activity | Cytoplasm |
| mfd cbac_03010 | WP_002518816.1 | Transcription-repair-coupling factor | Transcription | Binding | Cytoplasm |
| PPA1711 cbac_09310 | WP_002522145.1 | Short chain dehydrogenase | Lipid metabolism | Oxidoreductase activity | Cytoplasm |
| PPA1048 cbac_05685 | WP_002515631.1 | PglZ domain-containing protein | Cellular homeostasis | Catalytic activity | Cytoplasm |
| cbac_01755 | WP_002518719.1 | Sugar phosphate isomerase/epimerase | Metabolic process | Isomerase activity | Cytoplasm |
| cbac_02605 | WP_002518777.1 | Tat pathway signal protein | Transport | Transolcase activity | Membrane |
| cbac_12735 | WP_002518540.1 | Chromosome partitioning protein ParB | DNA replication | Binding | Cytoplasm |
| dnaA HMPREF0675_3000 | WP_002515747.1 | Chromosomal replication initiator protein DnaA | DNA replication | Binding | Cytoplasm |
| rbpA PPA0820 cbac_04490 | WP_002513867.1 | RNA polymerase-binding protein RbpA | Transcription | Binding | Unknown |
| gatB HMPREF0675_4186 | WP_002516598.1 | Aspartyl/glutamyl-tRNA | Translation | Transferase activity | Cytoplasm |
| pgl PPA1565 cbac_08515 | WP_002521397.1 | 6-phosphogluconolactonase | Metabolic process | Phosphogluconolactonase activity | Unknown |
| PPA1083 cbac_05885 | WP_002523969.1 | Glycine cleavage system protein H | Metabolic process | Transferase activity | Membrane |
| ligA cbac_08830 | WP_002531030.1 | DNA ligase | DNA repair | Ligase activity | Cytoplasm |
| ligA cbac_08830 | WP_002531030.1 | DNA ligase | DNA replication | Ligase activity | Cytoplasm |
| purK PPA1702 cbac_09260 | WP_002531235.1 | N5-carboxyaminoimidazole ribonucleotide synthase | Biosynthetic process | Synthase activity | Unknown |
| PPA1031 cbac_05600 | WP_002515632.1 | RNA polymerase principal sigma factor HrdD | Transcription | Binding | Cytoplasm |
| whiA HMPREF0675_3880 | WP_002515546.1 | Probable cell division protein WhiA | Cell division | Binding | Cytoplasm |
| cbac_01795 | WP_002531193.1 | Histidine phosphatase family protein | Regulation of biological process | Isomerase activity | Cytoplasm |
| cbac_02850 | WP_002518803.1 | Molybdopterin-guanine dinucleotide biosynthesis protein | Biosynthetic process | Binding | Cytoplasm |
| trpC PPA1130 cbac_06135 | WP_002516614.1 | Indole-3-glycerol phosphate synthase | Biosynthetic process | Synthase activity | Cytoplasm |
| cbac_00405 | WP_002515702.1 | ABC transporter ATP-binding protein | Transport | Binding | Membrane |
| PPA1364 cbac_07445 | WP_002517908.1 | IclR family transcriptional regulator | Transcription | Molecular function regulator | Cytoplasm |
| PPA0655 cbac_03620 | WP_002515176.1 | Antitoxin HicB | Response to stimulus | Binding | Unknown |
| PPA0055 cbac_00300 | WP_002515807.1 | DNA-binding response regulator | Transcription | Molecular function regulator | Nucleoid |
| rsmI PPA0524 cbac_02910 | WP_002516667.1 | Ribosomal RNA small subunit methyltransferase I | Response to stimulus | Transferase activity | Cytoplasm |
| PPA1824 cbac_09925 | WP_009640388.1 | Glycosyl transferase | Biosynthetic process | Transferase activity | Cytoplasm |
| PPA2237 cbac_12170 | WP_002516470.1 | 1-acyl-sn-glycerol-3-phosphate acyltransferase | Biosynthetic process | Transferase activity | Cytoplasm |
| PPA1146 cbac_06210 | WP_011183804.1 | Beta-glucosidase | Metabolic process | Hydrolase activity | Cytoplasm |
| cbac_05620 | WP_002515456.1 | Alpha/beta hydrolase | Degradation | Hydrolase activity | Membrane |
| HMPREF0675_5088 | WP_002518085.1 | Phosphomannomutase | Biosynthetic process | Hydrolase activity | Cytoplasm |
| cbac_06220 | WP_041444150.1 | Permease IIC component | Transport | Transferase activity | Membrane |
| PPA0082 cbac_00450 | WP_002515718.1 | Hypothetical membrane associated protein | Transport | Transporter activity | Membrane |
| rnhA PPA1729 cbac_09405 | WP_002531248.1 | Ribonuclease H | Metabolic process | Binding | Cytoplasm |
| PPA1898 cbac_10295 | WP_002531304.1 | Exodeoxyribonuclease III | DNA repair | Ribonuclease activity | Cytoplasm |
| PPA1069 cbac_05795 | WP_002517916.1 | Amino acid permease | Transport | Transporter activity | Membrane |
| PPA2185 cbac_11850 | WP_002516309.1 | Acyl dehydratase | Metabolic process | Isomerase activity | Cytoplasm |

| | | | | | |
|---|---|---|---|---|---|
| cbac_09895 | WP_002525147.1 | Glyco_hydro_35 domain-containing protein | Metabolic process | Hydrolase activity | Cytoplasm |
| cbac_06995 | WP_002524911.1 | Peptide synthetase | Biosynthetic process | Synthase activity | Cytoplasm |
| uvrB cbac_04410 | WP_002518347.1 | UvrABC system protein B | Response to stimulus | Binding | Cytoplasm |
| PPA1118 cbac_06075 | WP_009640307.1 | Methionine synthase vitamin-B12 independent | Biosynthetic process | Transferase activity | Cytoplasm |
| rsmG cbac_12765 | WP_002518545.1 | Ribosomal RNA small subunit methyltransferase G | Translation | Transferase activity | Cytoplasm |
| PPA0120 cbac_00670 | WP_002530833.1 | EXLDI protein | Unknown | Unknown | Unknown |
| PPA0624 cbac_03465 | WP_002530925.1 | N-acetyltransferase | Metabolic process | Transferase activity | Cytoplasm |
| PPA1166 cbac_06315 | WP_002517766.1 | Bifunctional | Metabolic process | Hydrolase activity | Cytoplasm |
| PPA2037 cbac_11045 | WP_002514480.1 | MerR family DNA-binding transcriptional regulator | Transcription | Molecular function regulator | Cytoplasm |
| PPA1186 | WP_002513526.1 | PPK2 domain-containing protein | Metabolic process | Kinase activity | Cytoplasm |
| PPA1616 cbac_08775 | WP_002517026.1 | DNA helicase | DNA replication | Hydrolase activity | Cytoplasm |
| PPA0348 cbac_01935 | WP_002517076.1 | Conserved protein, putative glycine cleavage T-protein | Metabolic process | Unknown | Unknown |
| PPA1488 cbac_08115 | WP_002520231.1 | Arabinose operon protein AraM | Biosynthetic process | Oxidoreductase activity | Cytoplasm |
| PPA1524 cbac_08315 | WP_009640297.1 | 6-phosphogluconolactonase | Metabolic process | Phosphogluconolactonase activity | Cytoplasm |
| cbac_04925 | WP_002515532.1 | Isochorismate synthase | Biosynthetic process | Synthase activity | Cytoplasm |
| cobO PPA0437 cbac_02385 | WP_002515069.1 | Cob | Biosynthetic process | Transferase activity | Unknown |
| manA PPA0451 cbac_02480 | WP_002517436.1 | Mannose-6-phosphate isomerase | Metabolic process | Isomerase activity | Cytoplasm |
| PPA1085 cbac_05895 | WP_002513631.1 | MerR family transcriptional regulator | Transcription | Molecular function regulator | Cytoplasm |
| PPA2254 cbac_12250 | WP_002516466.1 | Amino acid permease | Transport | Transporter activity | Membrane |
| PPA0639 cbac_03540 | WP_002530928.1 | Glycogen synthase | Biosynthetic process | Transferase activity | Cytoplasm |
| PPA0177 cbac_00965 | WP_002530809.1 | APC family permease | Transport | Transporter activity | Membrane |
| cbac_12740 | WP_002518541.1 | ParA family protein | Cell division | Binding | Cytoplasm |
| PPA0685 cbac_03785 | WP_002515306.1 | Sialidase | Metabolic process | Sialidase activity | Cytoplasm |
| cobI PPA0420 cbac_02305 | WP_002531167.1 | Precorrin-2 C | Biosynthetic process | Transferase activity | Cytoplasm |
| PPA1464 cbac_07995 | WP_009640171.1 | Methylase | Translation | Transferase activity | Cytoplasm |
| nhaA HMPREF0675_3274 | WP_002524662.1 | Na | Cellular homeostasis | Transporter activity | Membrane |
| PPA2031 cbac_11015 | WP_002530684.1 | ABC transporter associated permease | Transport | Transporter activity | Membrane |
| mraY HMPREF9578_00542 | WP_002513927.1 | Phospho-N-acetylmuramoyl-pentapeptide-transferase | Cell wall organization and cell shape | Transferase activity | Membrane |
| HMPREF9578_00411 | WP_002521769.1 | Metallo-beta-lactamase domain protein | Regulation of biological process | Hydrolase activity | Cytoplasm |
| cbac_03830 | WP_002523706.1 | Cytochrome c oxidase assembly protein | Protein processing | Unknown | Membrane |
| PPA1365 cbac_07460 | WP_011183848.1 | CPBP family intramembrane metalloprotease | Protein processing | Peptidase activity | Membrane |
| PPA0557 cbac_03110 | WP_002515113.1 | Transporter | Transport | Transporter activity | Membrane |
| thiL PPA1357 cbac_07420 | WP_002515338.1 | Thiamine-monophosphate kinase | Biosynthetic process | Binding | Cytoplasm |
| cbac_11435 | WP_002530754.1 | Histidine kinase | Regulation of biological process | Kinase activity | Membrane |
| HMPREF0675_4955 | WP_002515985.1 | Enoyl-CoA hydratase/isomerase family protein | Unknown | Isomerase activity | Cytoplasm |
| tyrS cbac_07675 | WP_009640057.1 | Tyrosine--tRNA ligase | Translation | Ligase activity | Cytoplasm |
| thrC HMPREF0675_4320 | WP_002518907.1 | Threonine synthase | Biosynthetic process | Synthase activity | Cytoplasm |
| PPA1258 cbac_06795 | WP_002517921.1 | Homoserine dehydrogenase | Biosynthetic process | Dehydrogenase activity | Cytoplasm |
| ddl PPA1359 cbac_07430 | WP_002531315.1 | D-alanine--D-alanine ligase | Cell wall organization and cell shape | Ligase activity | Cytoplasm |
| PPA0109 cbac_00605 | WP_011183602.1 | Myosin-crossreactive antigen | Lipid metabolism | Binding | Cytoplasm |
| cbac_09270 | WP_002517608.1 | Peptidase E | Metabolic process | Peptidase activity | Cytoplasm |

| | | | | | |
|---|---|---|---|---|---|
| PPA2251 cbac_12235 | WP_002518145.1 | D-glycerate dehydrogenase | Metabolic process | Dehydrogenase activity | Cytoplasm |
| cbac_10845 | WP_002518064.1 | Dihydrodipicolinate synthase family protein | Biosynthetic process | Lyase activity | Cytoplasm |
| PPA1367 cbac_07470 | WP_002516498.1 | Thymidine phosphorylase | Metabolic process | Phosphorylase activity | Cytoplasm |
| purQ HMPREF9578_01524 | WP_002514428.1 | Phosphoribosylformylglycinamidine synthase subunit PurQ | Biosynthetic process | Synthase activity | Cytoplasm |
| PPA2322 cbac_12635 | WP_002517539.1 | Conserved L-arabinose operon protein, hydrolase | Transcription | Hydrolase activity | Cytoplasm |
| PPA1285 cbac_06985 | WP_002517793.1 | ATP-grasp domain-containing protein | Energy process | Binding | Cytoplasm |
| PPA2238 cbac_12175 | WP_002516449.1 | AraC family transcriptional regulator | Transcription | Molecular function regulator | Cytoplasm |
| PPA2125 cbac_11505 | WP_007401162.1 | Polyphosphate glucokinase/transcriptional regulator | Transcription | Kinase activity | Cytoplasm |
| PPA1905 cbac_10330 | WP_002531307.1 | 2-oxoacid:ferredoxin oxidoreductase subunit beta | Metabolic process | Catalytic activity | Cytoplasm |
| cobF PPA1920 cbac_10410 | WP_002524506.1 | Precorrin-6A synthase | Biosynthetic process | Synthase activity | Cytoplasm |
| PPA2093 cbac_11320 | WP_002530734.1 | Alpha-ketoacid dehydrogenase subunit beta | Response to stimulus | Dehydrogenase activity | Cytoplasm |
| PPA0806 cbac_04425 | WP_002550711.1 | Maleylpyruvate isomerase family mycothiol-dependent enzyme | Metabolic process | Isomerase activity | Cytoplasm |
| murE PPA0753 cbac_04150 | WP_002518243.1 | UDP-N-acetylmuramoyl-L-alanyl-D-glutamate--2,6-diaminopimelate ligase | Biosynthetic process | Ligase activity | Cytoplasm |
| PPA0396 | WP_002515101.1 | L-serine dehydratase | Metabolic process | Binding | Cytoplasm |
| PPA1067 cbac_05785 | WP_002520491.1 | Transcriptional regulator | Transcription | Molecular function regulator | Cytoplasm |
| PPA1719 cbac_09355 | WP_002514720.1 | Biotin carboxylase | Lipid metabolism | Ligase activity | Cytoplasm |
| rplK HMPREF0675_4942 | WP_002516066.1 | 50S ribosomal protein L11 | Translation | Structural molecule activity | Ribosome |
| PPA0081 cbac_00440 | WP_002517298.1 | Alpha-1,4 glucan phosphorylase | Metabolic process | Phosphorylase activity | Cytoplasm |
| cbac_02400 | WP_002517307.1 | Cobalt-precorrin-6A reductase | Biosynthetic process | Reductase activity | Cytoplasm |
| hemE cbac_01710 | WP_002531196.1 | Uroporphyrinogen decarboxylase | Biosynthetic process | Decarboxylase activity | Cytoplasm |
| PPA1442 cbac_07865 | WP_002514249.1 | UPF0109 protein PPA1442 | Unknown | Binding | Unknown |
| cbac_11690 | WP_002516303.1 | ABC transporter ATP-binding protein | Transport | Transporter activity | Membrane |
| pyrG HMPREF0675_4439 | WP_002517885.1 | CTP synthase | Metabolic process | Binding | Cytoplasm |
| cbac_05930 | WP_002551083.1 | Threonylcarbamoyl-AMP synthase | Biosynthetic process | Binding | Cytoplasm |
| PPA0722 cbac_03995 | WP_002521859.1 | Glucokinase | Metabolic process | Kinase activity | Cytoplasm |
| rplT HMPREF9578_02089 | WP_002514221.1 | 50S ribosomal protein L20 | Translation | Structural molecule activity | Ribosome |
| typA bipA PPA2003 cbac_10865 | WP_002516314.1 | 50S ribosomal subunit assembly factor BipA | Translation | Binding | Ribosome |
| PPA0131 cbac_00730 | WP_002515744.1 | Glycosyl transferase | Biosynthetic process | Transferase activity | Cytoplasm |
| pdhA cbac_11325 | WP_002530735.1 | Pyruvate dehydrogenase | Unknown | Oxidoreductase activity | Unknown |
| cbac_02450 | WP_007400895.1 | PTS sugar transporter | Transport | Transporter activity | Membrane |
| carB cbac_05435 | WP_002531412.1 | Carbamoyl-phosphate synthase large chain | Biosynthetic process | Synthase activity | Cytoplasm |
| PPA0286 cbac_01600 | WP_002531205.1 | Conserved protein, putative cell division inhibitor | Cell division | Catalytic activity | Cytoplasm |
| PPA0147 cbac_00815 | WP_002515854.1 | Glycosyltransferase family 4 protein | Unknown | Transferase activity | Cytoplasm |
| cbac_06440 | WP_002524945.1 | 30S ribosomal protein S13 | Translation | Structural molecule activity | Ribosome |
| HMPREF0675_5340 | WP_002516467.1 | ErfK/YbiS/YcfS/YnhG | Biosynthetic process | Transferase activity | Membrane |
| PPA1056 cbac_05725 | WP_011183789.1 | Conserved protein | Unknown | Binding | Unknown |
| sigA cbac_05605 | WP_002519002.1 | RNA polymerase sigma factor SigA | Transcription | Binding | Cytoplasm |
| ettA PPA1636 cbac_08880 | WP_002516980.1 | Energy-dependent translational throttle protein EttA | Translation | Transferase activity | Cytoplasm |
| rbsK PPA0018 cbac_00095 | WP_002515739.1 | Ribokinase | Metabolic process | Kinase activity | Cytoplasm |
| cbac_03165 | WP_002519222.1 | PhoH family protein | Unknown | Binding | Unknown |
| thiM PPA0885 | WP_002531443.1 | Hydroxyethylthiazole kinase | Biosynthetic process | Binding | Cytoplasm |

| | | | | | |
|---|---|---|---|---|---|
| cbac_03675 | WP_002519174.1 | Glutamine synthetase | Biosynthetic process | Ligase activity | Cytoplasm |
| polA cbac_04270 | WP_002523754.1 | DNA polymerase I | DNA repair | Binding | Nucleoid |
| polA cbac_04270 | WP_002523754.1 | DNA polymerase I | DNA replication | Binding | Nucleoid |
| PPA0945 cbac_05135 | WP_002526277.1 | Putative histidine kinase | Regulation of biological process | Kinase activity | Membrane |
| cbac_12525 | WP_002519762.1 | Serine/threonine protein kinase | Regulation of biological process | Kinase activity | Cytoplasm |
| PPA0281 cbac_01575 | WP_002517113.1 | Conserved membrane spanning protein | Transport | Transporter activity | Membrane |
| radA PPA0312 cbac_01740 | WP_002517527.1 | DNA repair protein RadA | DNA repair | Hydrolase activity | Cytoplasm |
| PPA0342 cbac_01905 | WP_002517143.1 | Conserved protein | Unknown | Hydrolase activity | Cytoplasm |
| PPA1639 cbac_08895 | WP_002524831.1 | ABC transporter | Transport | Transporter activity | Membrane |
| PPA2096 cbac_11340 | WP_009639878.1 | Molybdenum cofactor biosynthesis enzyme/coproporphyrinogen III oxidase | Biosynthetic process | Catalytic activity | Cytoplasm |
| cbac_04120 | WP_002518266.1 | DUF58 domain-containing protein | Unknown | Structural molecule activity | Membrane |
| PPA0710 cbac_03930 | WP_002530941.1 | Cytochrome bc1 complex cytochrome b subunit | Unknown | Reductase activity | Membrane |
| hrcA HMPREF9578_00410 | WP_002513807.1 | Heat-inducible transcription repressor HrcA | Response to stimulus | Binding | Cytoplasm |
| PPA0072 cbac_00395 | WP_002525540.1 | Membrane associated protein | Transport | Transporter activity | Membrane |
| PPA2298 cbac_12500 | WP_002515857.1 | Conserved protein, putative mechanosensitive ion channel | Transport | Transporter activity | Membrane |
| PPA0732 cbac_04045 | WP_002513952.1 | Geranylgeranyl pyrophosphate synthase | Biosynthetic process | Transferase activity | Cytoplasm |
| PPA1671 cbac_09095 | WP_002522165.1 | Dihydrofolate reductase | Metabolic process | Reductase activity | Cytoplasm |
| PPA1917 cbac_10395 | WP_002515372.1 | Hypothetical membrane protein | Transport | Transporter activity | Membrane |
| PPA0434 | WP_002517347.1 | ABC transporter ATP-binding protein | Transport | Transferase activity | Membrane |
| PPA1476 cbac_08060 | WP_002516879.1 | Glycine betaine transport system permease protein | Transport | Transporter activity | Membrane |
| PPA1955 cbac_10590 | WP_002524492.1 | Right-handed parallel beta-helix repeat-containing protein | Unknown | Catalytic activity | Unknown |
| cbac_04030 | WP_002521854.1 | Non-specific serine/threonine protein kinase | Regulation of biological process | Kinase activity | Cytoplasm |
| nuoH HMPREF9578_02182 | WP_002514928.1 | NADH-quinone oxidoreductase subunit H | Unknown | Oxidoreductase activity | Membrane |
| cbac_00710 | WP_002526015.1 | DUF2029 domain-containing protein | Unknown | Structural molecule activity | Membrane |
| PPA0128 cbac_00715 | WP_011183607.1 | Membrane spanning protein | Biosynthetic process | Transferase activity | Membrane |
| PPA2033 cbac_11025 | WP_002530685.1 | Amino acid permease | Transport | Transporter activity | Membrane |

#### Table S2. List of proteins identified in *C. acnes*-derived H1 EVs under anaerobic conditions

| Gene name | Acession number | Protein name | GO - Biological process | GO - Molecular function | GO - Cellular component |
|---|---|---|---|---|---|
| HMPREF9578_00571 | WP_002521853.1 | Ribonuclease | Cellular homeostasis | Ribonuclease activity | Cytoplasm |
| pdxA cbac_01665 | WP_002531202.1 | 4-hydroxythreonine-4-phosphate dehydrogenase PdxA | Biosynthetic process | Oxidoreductase activity | Cytoplasm |
| folE PPA0261 cbac_01480 | WP_002517093.1 | GTP cyclohydrolase 1 | Metabolic process | Binding | Cytoplasm |
| PPA0503 cbac_02790 | WP_002518798.1 | Helix-turn-helix domain-containing protein | Regulation of biological process | Binding | Unknown |
| mmsA PPA0461 cbac_02540 | WP_002531156.1 | Methylmalonate-semialdehyde dehydrogenase | Metabolic process | Dehydrogenase activity | Cytoplasm |
| PPA1815 cbac_09865 | WP_002517606.1 | 3-oxoacyl-ACP reductase FabG | Lipid metabolism | Reductase activity | Cytoplasm |
| PPA1609 cbac_08735 | WP_002531040.1 | Conserved phage-associated protein | Unknown | Unknown | Unknown |
| PPA2063 cbac_11190 | WP_002530709.1 | Peptide ABC transporter substrate-binding protein | Transport | Transporter activity | Membrane |
| PPA0851 cbac_04650 | WP_002531018.1 | ATP-dependent endonuclease | DNA repair | Hydrolase activity | Cytoplasm |
| PPA0144 cbac_00800 | WP_011183610.1 | DeoR/GlpR transcriptional regulator | Transcription | Molecular function regulator | Nucleoid |
| PPA1638 cbac_08890 | WP_002531028.1 | ABC transporter | Transport | Transporter activity | Membrane |
| PPA2091 cbac_11310 | WP_002530732.1 | ABC transporter ATP-binding protein | Transport | Transporter activity | Membrane |
| PPA1760 cbac_09575 | WP_002514763.1 | ABC transporter associated permease | Transport | Transporter activity | Membrane |
| PPA0399 cbac_02205 | WP_002531171.1 | Sugar ABC transporter | Transport | Transporter activity | Membrane |
| PPA1147 cbac_06215 | WP_002513567.1 | PTS sugar transporter subunit IIB | Transport | Kinase activity | Membrane |
| PPA1915 cbac_10385 | WP_002518040.1 | ABC transporter ATP-binding protein | Energy process | Binding | Membrane |
| cbac_02210 | WP_002517551.1 | Carbohydrate ABC transporter permease | Transport | Transporter activity | Membrane |
| PPA0925 cbac_05045 | WP_002513798.1 | Putative two-component system sensor kinase | Response to stimulus | Kinase activity | Membrane |
| PPA1599 cbac_08685 | WP_007400937.1 | CAAX protease | Protein processing | Peptidase activity | Membrane |
| PPA1389 cbac_07600 | WP_002516495.1 | NUDIX hydrolase | Cellular homeostasis | Hydrolase activity | Cytoplasm |
| PPA1721 cbac_09365 | WP_002522139.1 | Dihydrolipoamide dehydrogenase | Cellular homeostasis | Dehydrogenase activity | Cytoplasm |
| iolB PPA0458 cbac_02525 | WP_002531157.1 | 5-deoxy-glucuronate isomerase | Metabolic process | Isomerase activity | Cytoplasm |
| PPA0398 cbac_02200 | WP_002520024.1 | Conserved protein, putative sugar-binding protein | Regulation of biological process | Binding | Membrane |
| cbac_10405 | WP_002519594.1 | CoA ester lyase | Unknown | Lyase activity | Unknown |
| PPA1529 cbac_08340 | WP_002516971.1 | Conserved protein | Unknown | Unknown | Membrane |
| moaC PPA0498 cbac_02765 | WP_007401059.1 | Cyclic pyranopterin monophosphate synthase | Biosynthetic process | Synthase activity | Cytoplasm |
| menB PPA0907 cbac_04940 | WP_002513816.1 | 1,4-dihydroxy-2-naphthoyl-CoA synthase | Biosynthetic process | Synthase activity | Cytoplasm |
| PPA1229 cbac_06650 | WP_002517946.1 | ArsR family transcriptional regulator | Transcription | Molecular function regulator | Cytoplasm |
| PPA1086 cbac_05900 | WP_002517877.1 | Bifunctional nuclease family protein | Protein processing | Hydrolase activity | Nucleoid |
| PPA2112 cbac_11430 | WP_002530753.1 | DNA-binding response regulator | Transcription | Binding | Nucleoid |
| PPA1459 cbac_07975 | WP_002514269.1 | Alpha-mannosidase | Metabolic process | Binding | Cytoplasm |
| PPA1050 cbac_05695 | WP_011183787.1 | Conserved protein | Unknown | Unknown | Unknown |
| PPA0062 cbac_00345 | WP_002530850.1 | Alpha-mannosidase | Metabolic process | Binding | Cytoplasm |
| aceE cbac_05385 | WP_002525003.1 | Pyruvate dehydrogenase E1 component | Metabolic process | Dehydrogenase activity | Cytoplasm |
| PPA2267 cbac_12320 | WP_002519750.1 | 6-phosphofructokinase | Metabolic process | Kinase activity | Cytoplasm |
| topA PPA0241 cbac_01375 | WP_002529762.1 | DNA topoisomerase 1 | DNA replication | Binding | Nucleoid |
| nadE PPA2266 cbac_12310 | WP_002518158.1 | Glutamine-dependent NAD | Biosynthetic process | Synthase activity | Cytoplasm |
| PPA1142 cbac_06190 | WP_002521618.1 | PTS system, sucrose-specific IIBC component | Transport | Transferase activity | Membrane |
| cbac_04110 | WP_002530961.1 | DUF3040 domain-containing protein | Unknown | Unknown | Membrane |
| aroC cbac_06400 | WP_002517990.1 | Chorismate synthase | Biosynthetic process | Synthase activity | Cytoplasm |

| | | | | | |
|---|---|---|---|---|---|
| cbac_10960 | WP_002519475.1 | MFS transporter | Transport | Transporter activity | Membrane |
| gnpA PPA0083 cbac_00455 | WP_002530846.1 | 1,3-beta-galactosyl-N-acetylhexosamine phosphorylase | Metabolic process | Phosphorylase activity | Cytoplasm |
| PPA2320 cbac_12625 | WP_002517435.1 | L-arabinose utilization protein, glycerol dehydrogenase | Lipid metabolism | Oxidoreductase activity | Cytoplasm |
| PPA0290 cbac_01620 | WP_002517197.1 | Putative two-component sensor kinase | Response to stimulus | Structural molecular activity | Membrane |
| ctaD PPA0702 cbac_03885 | WP_002518277.1 | Cytochrome c oxidase subunit 1 | Transport | Binding | Membrane |
| PPA2203 cbac_11960 | WP_002518166.1 | Putative Na+/H+ antiporter | Transport | Transporter activity | Membrane |
| PPA1315 cbac_07150 | WP_009640225.1 | Exodeoxyribonuclease V gamma chain | DNA repair | Ribonuclease activity | Cytoplasm |
| cbac_10755 | WP_002523368.1 | Pyridine nucleotide-disulfide oxidoreductase | Cellular homeostasis | Oxidoreductase activity | Nucleoid |
| PPA1610 cbac_08740 | WP_002531039.1 | Putative transcriptional regulator | Transcription | Binding | Nucleoid |
| PPA0416 cbac_02280 | WP_011183669.1 | DNA-binding response regulator | Transcription | Binding | Nucleoid |
| PPA2216 cbac_12040 | WP_002514161.1 | Acyl-CoA dehydrogenase | Lipid metabolism | Dehydrogenase activity | Cytoplasm |
| PPA1273 cbac_06890 | WP_002516865.1 | ABC transporter ATP-binding protein | Transport | Transporter activity | Membrane |
| cbac_02265 | WP_002517453.1 | ABC transporter ATP-binding protein | Transport | Transporter activity | Membrane |
| cobA PPA0439 cbac_02395 | WP_002517252.1 | Uroporphyrinogen III methylase | Biosynthetic process | Transferase activity | Cytoplasm |
| argC HMPREF0675_4395 | WP_002520173.1 | N-acetyl-gamma-glutamyl-phosphate reductase | Biosynthetic process | Reductase activity | Cytoplasm |
| PPA2088 cbac_11295 | WP_002530729.1 | Esc family protein | Unknown | Unknown | Membrane |
| rsmH cbac_04135 | WP_002530963.1 | Ribosomal RNA small subunit methyltransferase H | Translation | Transferase activity | Cytoplasm |
| PPA1217 cbac_06590 | WP_002518927.1 | 4-hydroxybutyrate coenzyme A transferase | Metabolic process | Transferase activity | Cytoplasm |
| PPA0141 cbac_00780 | WP_002530823.1 | HPr family phosphocarrier protein | Transport | Transferase activity | Cytoplasm |
| PPA2240 cbac_12185 | WP_002530896.1 | Cation-transporting ATPase | Transport | Transporter activity | Membrane |
| HMPREF0675_5099 | WP_002516194.1 | TrkA C-terminal domain protein | Transport | Transporter activity | Membrane |
| PPA1732 cbac_09420 | WP_002531250.1 | ABC transporter associated permease | Transport | Transporter activity | Membrane |
| PPA0540 cbac_03005 | WP_002516643.1 | Conserved protein // PrsW family intermembrane metalloprotease | Regulation of biological process | Peptidase activity | Membrane |
| PPA0068 cbac_00375 | WP_002519827.1 | Putative two component sensor kinase | Response to stimulus | Kinase activity | Membrane |
| PPA1275 cbac_06900 | WP_002516777.1 | ABC transporter permease | Transport | Transporter activity | Membrane |

Universitat Pompeu Fabra Barcelona

#### Table S3. List of proteins identified in *C. acnes*-derived H2 EVs under anaerobic conditions

| Gene name | Accession number | Protein name | GO - Biological Process | GO - Molecular function | GO - Cellular component |
|---|---|---|---|---|---|
| PPA1345 cbac_07350 | WP_002517702.1 | DUF1707 domain-containing protein | Unknown | Unknown | Membrane |
| HMPREF0675_3894 | WP_002515660.1 | Uncharacterized protein | Unknown | Unknown | Membrane |
| rpsT HMPREF0675_3956 | WP_002515457.1 | 30S ribosomal protein S20 | Translation | Structural molecular activity | Ribosome |
| PPA0522 cbac_02900 | WP_011183691.1 | Nitrate reductase | Energy process | Transporter activity | Membrane |
| PPA0960 cbac_05215 | WP_002517830.1 | MsnO8 family LLM class oxidoreductase | Regulation of biological process | Oxidoreductase activity | Membrane |
| PPA0960 cbac_05215 | WP_002517830.1 | MsnO8 family LLM class oxidoreductase | Response to stimulus | Oxidoreductase activity | Cytoplasm |
| PPA0724 cbac_04005 | WP_002530950.1 | 1-acyl-sn-glycerol-3-phosphate acyltransferase | Lipid metabolism | Transferase activity | Cytoplasm |
| hisB HMPREF9578_00150 | WP_002513559.1 | Imidazoleglycerol-phosphate dehydratase | Byosynthetic process | Lyase activity | Cytoplasm |
| kup cbac_11865 | WP_002519550.1 | Probable potassium transport system protein kup | Transport | Transporter activity | Membrane |
| uppP HMPREF0675_5297 | WP_002516473.1 | Undecaprenyl-diphosphatase | Cell wall organization and cell shape | Phosphatase activity | Membrane |
| PPA0619 cbac_03440 | WP_002530923.1 | Allantoin permease | Transport | Transporter activity | Membrane |

Universitat Pompeu Fabra Barcelona

### Table S4. List of primer sequences used for RT-qPCR

| Genes | | Primers (From 5' to 3') |
|---|---|---|
| CREBBP | FWD | 5' GAGAGCAAGCAAACGGAGAG |
| | RV | 5' AAGGGAGGCAAACAGGACA |
| TNFα SEB | FWD | 5' CCAGGGACCTCTCTCTAATCA |
| | RV | 5' TCAGCTTGAGGGTTTGCTAC |
| IL-6 SEB | FWD | 5' ACTCACCTCTTCAGAACGAATTG |
| | RV | 5' AGCCATCTTTGGAAGGTTCAG |
| IL-8 SEB | FWD | 5' CTTGGCAGCCTTCCTGATTT |
| | RV | 5' GGGTGGAAAGGTTTGGAGTATG |
| PLIN 2 | FWD | 5' TCAGCTCCATTCTACTGTTCACC |
| | RV | 5' CCTGAATTTTCTGATTGGCACT |
| TGFβ1 | FWD | 5' TACCTGAACCCGTGTTGCTCTC |
| | RV | 5' GTTGCTGAGGTATCGCCAGGAA |
| COX-2 | FWD | 5' GAATCATTCACCAGGCAAATTG |
| | RV | 5' TCTGTACTGCGGGTGGAACA |
| OCLN | FWD | 5' GTCATCCAGGCCTCTTGAAA |
| | RV | 5' GGTGATAATGATTCGGTTTG |
| MMP2 | FWD | 5' AGCGAGTGGATGCCGCCTT |
| | RV | 5' CATTCCAGGCATCTGCGAT |
| CLDN1 | FWD | 5' GTCTTTGACTCCTTGCTGAATCTG |
| | RV | 5' CACCTCATCGTCTTCCAAGCAC |
| IL-10 | FWD | 5' TCTCCGAGATGCCTTCAGCAGA |
| | RV | 5' TCAGACAAGGCTTGGCAACCCA |
